# Multi-Omics Regulatory Network Inference in the Presence of Missing Data

**DOI:** 10.1101/2022.04.14.488153

**Authors:** Juan D. Henao, Michael Lauber, Manuel Azevedo, Anastasiia Grekova, Fabian Theis, Markus List, Christoph Ogris, Benjamin Schubert

## Abstract

A key problem in systems biology is the discovery of regulatory mechanisms that drive phenotypic behaviour of complex biological systems in the form of multi-level networks. Modern multi-omics profiling techniques probe these fundamental regulatory networks but are often hampered by experimental restrictions leading to missing data or partially measured omics types for subsets of individuals due to cost restrictions. In such scenarios, in which missing data is present, classical computational approaches to infer regulatory networks are limited. In recent years, approaches have been proposed to infer sparse regression models in the presence of missing information. Nevertheless, these methods have not been adopted for regulatory network inference yet.

In this study, we integrated regression-based methods that can handle missingness into KiMONo, a **K**nowledge gu**I**ded **M**ulti-**O**mics **N**etw**o**rk inference approach, and benchmarked their performance on commonly encountered missing data scenarios in single- and multi-omics studies. Overall, two-step approaches that explicitly handle missingness performed best for a wide range of random- and block-missingness scenarios on imbalanced omics-layers dimensions, while methods implicitly handling missingness performed best on balanced omics-layers dimensions. Our results show that robust multi-omics network inference in the presence of missing data with KiMONo is feasible and thus allows users to leverage available multi-omics data to its full extent.

**Juan Henao** is a 3rd year PhD candidate at Computational Health Center at Helmholtz Center Munich working on multi-omics and clinical data integration using both, bulk and single-cell data.

**Michael Lauber** is a PhD Candidate at the Chair of Experimental Bioinformatics at the Technical University Munich. Currently, he is working on an approach for inference of reprogramming transcription factors for trans-differentiation.

**Manuel Azevedo** is a Master’s student at the Technical University of Munich in Mathematics with a focus on Biomathematics and Biostatistics. Currently, he is working as a Student Assistant at Helmholtz Munich, where he is also doing his master’s thesis.

**Anastasiia Grekova** is a Master’s student of bioinformatics at the Technical University of Munich and the Ludwig-Maximilians-University Munich, working on multi-omics data integration in Marsico Lab at HMGU.

**Fabian Theis** is the Head of the Institute of Computational Biology and leading the group for Machine Learning at Helmholtz Center Munich. He also holds the chair of ‘Mathematical modelling of biological systems’, Department of Mathematics, Technical University of Munich as an Associate Professor.

**Markus List** obtained his PhD at the University of Southern Denmark and worked as a postdoctoral fellow at the Max Planck Institute for Informatics before starting his group Big Data in BioMedicine at the Technical University of Munich.

**Christoph Ogris** holds a PostDoc position in the Marsico Lab at Helmholtz-Center Munich. His research focuses on predicting and exploiting multi-modal biological networks to identify disease-specific cross-omic interactions.

**Benjamin Schubert** obtained his PhD at the University of Tübingen and worked as a postdoctoral fellow at Harvard Medical School and Dana-Farber Cancer Institute USA before starting his group for Translational Immmunomics at the Helmholtz Center Munich.

## Introduction

Complex biological systems are organised in multi-level, dynamically controlled networks that regulate and maintain the phenotypic behaviour of individual cells and their response to environmental changes [1]. Uncovering these multi-level networks and systemically understanding the interplay of their elements is a key problem in computational biology. Modern high-throughput multi-omics techniques now enable access to each regulatory network level, even at single-cell resolution [2,3]. However, combining multi-omics measurements and reconstructing the underlying regulatory network remains challenging [4]. Generally, sparse interaction networks in the form of directed or undirected graphs are constructed from dynamic interventional omics or large observational data using different classes of statistical methods [4]. Common approaches are either correlation-based [5], use techniques from information theory [6–8], or use (regularised) regression and variable selection frameworks to infer graphical models [9–11]. Most recent methods also integrate prior knowledge [12,13], such as experimentally determined protein-protein interaction networks, known metabolic pathways, or even predicted miRNA-mRNA interactions [14].

One such recent approach is KiMONo, **K**nowledge gu**i**ded **M**ulti-**O**mics **N**etw**o**rk inference [15], a two-step prior knowledge-based approach for multi-omics regulatory network inference. In the first step, the framework uses the whole dataset to model each omics element individually, detecting statistical effects between them. The framework combines all models in a second step, assembling a multi-omics graph with the input features as nodes linked via edges representing the detected effects.

However, a major drawback of most network inference methods is their inability to handle missing data. Many omics technologies such as mass-spectrometry-based proteomics and metabolomics or single-cell transcriptomics suffer from inherent missingness due to the stochasticity of the biological processes and technical limitations. Additionally, it is often necessary to combine multiple studies that only partially measure the same omics levels to reach sample sizes adequate for network inference, creating patterns of block-wise missingness. Many classical regulatory network inference methods ignore missing data and focus only on analysing complete cases, thus underutilising the collected data and severely limiting the amount of information used. Removing samples with missing features can also lead to biased estimates if the missingness is not completely random [16], potentially affecting the extracted regulatory network. Multiple imputation [17] is another popular approach to deal with missingness, followed by applying any classical network inference method using *ad hoc* rules to harmonise variable selection across multiply-imputed datasets [18]. However, Ganti and Willet demonstrated that such two-step approaches can be sub-optimal [19] and instead require integrated or more general frameworks to handle missing data and variable selection jointly.

In recent years, advances have been made in using sparse graphical models for data with missing information. These approaches can be roughly categorised in Bayesian methods using data augmentation strategies [20], methods using pooled posterior [21], bootstrapped inclusion probabilities [22,23], methods performing variable selection through stacked [18,24] or group Lasso integrated multiple imputation methods [25–28], low-rank matrix completion [19,29], inverse probability weighting [30], Lasso regularised inverse covariance estimation [31–34], and Expectation-Maximisation-based approaches [35,36]. While most methods address the missingness of individual features, some methods exist that also explicitly model block-missingness [28,37–39].

Incorporating such approaches in multi-omics network inference is attractive. However, a comprehensive benchmark of existing methods that can handle missing data is lacking. We, therefore, extended KiMONo with various regression-based approaches that integrate and combine prior imputed data [28] and Lasso-regularised inverse covariance estimation methods [33,34]. We systematically evaluated how these methods handle gradually increasing levels of artificial noise and missingness for regulatory network inference on single- and multi-omics data.

We observed that approaches explicitly handling missingness in a two-step manner performed best over a wide range of random, block-missingness, and noise levels in an imbalanced multi-omics dimensional dataset (TCGA-BRCA), while implicit covariance-estimation-based methods performed best in multi-omics with balanced dimensional dataset (TCGA-MIBC and TCGA-PRAD).

## Methods

### Knowledge guIded Multi-Omics Network (KiMONo) inference

KiMONo, **K**nowledge gu**I**ded **M**ulti-**O**mics **N**etw**o**rk inference [15], is a two-step inference procedure to detect statistical dependencies between omics features and construct a multi-omics network. Here, the inference complexity is decreased by pre-selecting feature dependencies based on existing prior knowledge of biological mechanisms such as known protein-protein interaction. Based on such prior knowledge, a system of linear multivariate regression models with sparse-group Lasso penalty is constructed [40]:

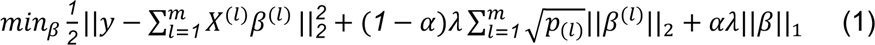

where α = [*0*,*1*], and *l* the group index of *m* groups (omics layer), being *X*^(*l*)^, *β*^(*l*)^, *p*_(*l*)_ the submatrix, coefficients, and length of coefficients for group *l*, respectively. This allows two types of sparsities: (1) “groupwise sparsity”, referring to all groups with at least one non-zero coefficient, and (2) “within group sparsity”, referring to all non-zero coefficients within every specific group (omics layer). The framework combines all models in a second step, assembling a multi-omics graph connecting regressed omics elements with its non-zero regressor elements if the regression performance (coefficient of determination) exceeds a predefined threshold.

### Regression-based Methods for Network Inference and Imputation

We focused on methods with a working R implementation and consistent documentation. These requirements left us with five advanced statistical approaches of three categories (1) stacked and (2) grouped multiple-imputation, as well as (3) Lasso-based inverse covariance estimation approaches (Table 1). All mentioned methods have been integrated into the KiMONo framework (https://github.com/cellmapslab/kimono). The data underlying this article, including detailed benchmarking results and code are available at Zenodo [41] (https://doi.org/10.5281/zenodo.7900595).

**Table 1:** Inference models included in this benchmark and capable of dealing with missing data.

| Method | Category | Citation |
| --- | --- | --- |
| knnKiMONo | single imputation + KiMONo | (Troyanskaya *et al.*, 2001);(Ogris *et al.*, 2021) |
| S(A)Lasso | stacked multiple imputation | (Du *et al.*, 2022) |
| G(A)Lasso | grouped multiple imputation | (Du *et al.*, 2022) |
| HMLasso | inverse covariance estimation | (Takada *et al.*, 2018) |
| CoCoLasso | inverse covariance estimation | (Datta and Zou, 2017);<br>(Takada *et al.*, 2018) |
| BDCoCoLasso | inverse covariance estimation | (Escribe *et al.*, 2021) |

#### kNN-imputation & Sparse Group Lasso (knnSGLasso)

We implemented a two-step approach that first imputes missing information using nearest neighbour averaging followed by applying the Sparse Group Lasso approach of the classical KiMONo. The kNN-based imputation method [42] implemented in the R package impute v1.46.0 was applied separately to individual omics layers and other covariates. Originally designed for the imputation of gene expression data, the method replaces missing values by averaging non-missing values of its nearest neighbours. If the percentage of missing data allowed for every variable (e.g., 50% per gene (default)) is exceeded, the missing values are imputed using the overall mean per sample. Only samples with missingness less than 80% (default) were considered for the imputation. The algorithm’s parameters were set to default values: the number of neighbours used in the imputation was set to k=10, and the largest block of variables imputed using the kNN algorithm before recursively dividing the feature into smaller chunks was set to max = 1500.

#### Stacked Adaptive LASSO (SALasso)

Stacked approaches combine prior *D*-times multiply imputed datasets by averaging over them, making such approaches applicable to existing sparse regression frameworks:

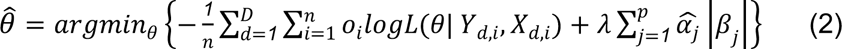

This results in a pooled estimate 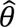 across all imputed datasets, thereby enforcing uniform variable selection across *p* covariates. Here α_*j*_ are so-called adaptive weights used to address estimation and selection inconsistencies that can occur and are defined as 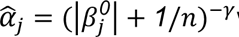 with some *γ* > 0 and 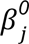 initial estimates obtained with (stacked)-Lasso or (stacked)-elastic-net. Following Du et al., *γ* is set to 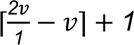 [28]. As stacking multiple imputed datasets can be seen as artificially increasing the sample size thereby potentially creating oversampling biases, observation weights *o_i_* for each subject can be included to account for the artificial inflation of the dataset with either uniform weights or with weights accounting for varying degrees of missingness per subject. We used the SALasso implementation released in the R package miselect 0.9.0 [28]. We tested 50 λ values with a lambda.min.ratio of 1e-4 in a 5-fold cross-validation with and without adaptive weights (SLasso), while the sample weights *o_i_* were set to be uniform.

#### Grouped Adaptive LASSO (GALasso)

Similar to SALasso, GALasso pools across prior imputed datasets by adding a group Lasso penalty term enforcing consistent variable selection across multiply imputed datasets yielding:

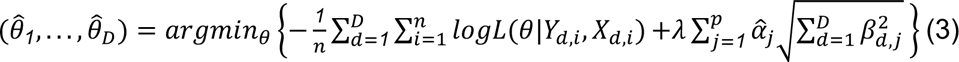

with 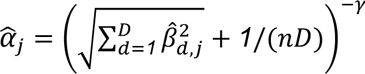, where 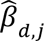 is the estimate obtained with 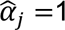, λ = ⌈*2v*/*1* − *v*⌉ + *1*, and *v* = *log*(*pD*)/*log*(*nD*) [28]. Contrary to SALasso, individual sets of parameters *θ_d_* are obtained for each of the *D* imputed datasets. However, the group-Lasso regularisation term jointly shrinks all *β*_.,j_ to zero, despite their potentially different estimates across the *D*-imputed datasets enabling a uniform variable selection. We used the GALasso implementation released in the R package miselect 0.9.0 [28]. We tested 50 λ values with a lambda.min.ratio of 1e-4 in a 5-fold cross-validation with and without adaptive weights (GLasso).

#### Convex Conditioned Lasso (CoCoLasso)

CoCoLasso is an inverse covariance estimation method for high-dimensional data with missing values. The main idea is to reformulate the Lasso regression by working with the sample covariance matrix of *X*, 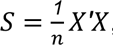, and the sample covariance vector of X and y, 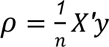. With this reformulation *β* is estimated via *S* and *ρ* instead of *X* and *y*. Since we are dealing with missing data, CoCoLasso works with the pairwise covariance *S^pair^* : = (*S_jk_^pair^*) and *ρ^pair^* : = (*ρ_jk_^pair^*) instead of *S* and *ρ*, where

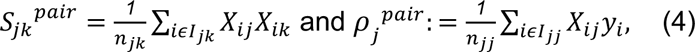

where *I_jk_*: = {*i*: *X_ij_* and *X_ik_* are observed}, and *n_jk_* is the number of elements of *I_jk_*. Then the inference problem can be defined as:

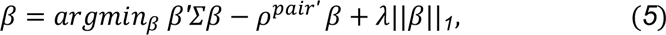

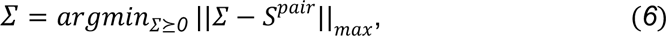

Since the matrix *S^pair^* needs to be positive semidefinite, CoCoLasso applies in (5) the alternating direction method of multipliers algorithm to obtain a positive semidefinite covariance matrix, *Σ*. Afterwards, CoCoLasso optimises the Lasso objective function (4). We used the CoCoLasso implementation released in the R package HMLasso 0.0.1 [34] with the following selection of hyperparameters: For λ, we tested 50 values with a lambda.min.ratio of 1e-1 in a 5-fold cross-validation.

To improve the computational efficiency of CoCoLasso, BDCoCoLasso was developed by implementing a block coordinate descent strategy [43] over corrupted (with missing data) and uncorrupted (full data) covariates. However, our preliminary results showed the random missingness of 5% and more encompassed already all samples and therefore BDCoCoLasso could not be applied. We, thus, refrain from discussing the performance of BDCoCoLasso.

#### Lasso with High Missing rate (HMLasso)

HMLasso can be seen as an optimally weighted modification of CoCoLasso according to the missing ratio. HMLasso uses the mean imputation method. Instead of *X*, the mean imputed data variable, *Z* is used, where *Z*_jk_ = *X_jk_* for an observed element and *Z*_jk_ = *0* otherwise. Followed by that the *S*^pair^ covariance matrix from CoCoLasso is modified by the covariance matrix of the mean imputed data, defined as *S^imp^* = *RS*^pair^, where *R*_jk_ = *n_jk_/n*.

The HMLasso is then formulated as:

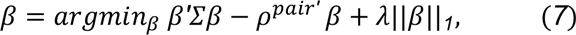

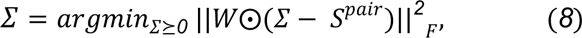

Where *W* = *R_jk_^α^* is defined as the weight matrix and α is the weight power parameter. We obtain the positive semi-definite matrix by minimising the weighted Frobenius norm in (8). Afterwards, HMLasso optimises the Lasso objective (7). Note that if we set α = *0* (non-weighted case), it is equal to CoCoLasso, replacing the Frobenius norm with the max norm. We used the HMLasso implementation released in the R package HMLasso 0.0.1 [34] with the following selection of hyperparameters: For α, we tested values between 0.5 and 2 with an interval of 0.5. For λ, we tested 50 values with a lambda.min.ratio of 1e-1 in a 5-fold cross-validation.

### Datasets

For the benchmarking performance evaluation, we collected two datasets consisting of triple-omics data (transcriptomics, copy number variation (CNV), and methylation) from the PanCancer Projects [44] using The Cancer Genome Atlas (TCGA) data portal and the cBioPortal [45]. Both datasets were already pre-processed including normalised and log-transformed gene expression, beta-values for methylation, and linear CNVs.

The first dataset encompassed 871 patients with breast invasive carcinoma (retrieved on 03/07/2022), while the second dataset comprised 414 patients with muscle-invasive bladder cancer (retrieved on 23/08/2022). All samples containing missing information were removed to construct a complete data set as the baseline, thus restricting the data sets to 604 and 404 samples, respectively. Similarly, features with low variance were removed, resulting in 11,530 transcriptomics, 1,366 methylation, and 84 CNV features for the first dataset (TCGA-BRCA from now on), and 20,085 transcriptomics, 15,460 methylation, and 24,765 CNV features for the second dataset (TCGA-MIBC from now on), respectively. These two datasets allowed us to study the impact of data dimensionality on performance.

A third dataset was collected from the PanCancer Projects [44] using The Cancer Genome Atlas (TCGA) data portal and the cBioPortal [45] for prostate adenocarcinoma (TCGA-PRAD from now on) (retrieved on 02/03/2023) comparable to TCGA-MIBC to evaluate performance consistency across similar datasets. The third dataset encompassed 491 samples with 20,123 transcriptomics, 15,576 methylation, and 24,765 CNV features after low variance removal.

### Prior network generation

For the first dataset (TCGA-BRCA), we used the prior network of the original KiMONo publication [15]. Briefly, protein-protein interactions were extracted from the BioGrid interactome (Release 3.5.188) [46], associated with the extracted gene expression information, and linked each CNV and methylation site to its associated gene since both omics layers were already annotated to gene identifiers. The final prior network contained 11,645 nodes (10,848 genes, 84 CNVs, and 713 methylation sites).

The construction of the prior network for the second and third datasets (TCGA-MIBC and TCGA-PRCA) followed the same steps as the first prior network construction using FunCoup v5 [47] (built-in September 2020) as the basis. We used interactions with maximum Final Bayesian Score (FBS) supporting protein-protein interactions, and log-likelihood ratio for physical protein-protein interaction evidence ≥ 99 percentile to extract reliable protein-protein interactions. The final prior network contained 51,318 nodes (18,990 genes, 18,460 CNVs, and 13,868 methylation sites).

### Network-based multiple imputation

Stacked and grouped adaptive Lasso approaches require multiple imputed data as input. However, multiple imputation methods do not scale well to high-dimensional data with high missingness, and standard implementations such as those offered in the R package MICE, take multiple hours to days to finish. Thus, we developed a novel network-guided multiple imputation by chained equation approach (ngMICE) by utilising KiMONo’s prior network. Instead of considering all covariates for imputation, we restrict each imputation attempt to the covariates that are directly linked to the missing covariate in a prior network as other covariates will be removed during network inference by KiMONo and therefore can be neglected. The number of covariates can be further reduced by correlation-based filtering. For missing elements retaining less than *k* covariates for imputation, the top *k* correlated covariates are used. Once the covariate matrix has been constructed as described, the standard MICE procedure is run. We used the R package MICE 3.14.0 [48], with predictive mean matching (default) as a multiple imputation approach, an absolute Pearson correlation coefficient of

0.1 as the threshold, and k = 5 most correlated features in the final regression formula. ngMICE performed similarly to kNN-based imputation in terms of RMSE across omics types and missingness with slightly worse average performance (Supplementary Figure 1).

### Benchmark

We assessed the performance of the selected inference methods in the presence of missing data by simulating three typical scenarios - (1) random missing information in a single omics level and across (2) multiple levels, as well as (3) block-missingness structures (Figure 1a). To test the methods’ capabilities even further, we decreased the signal-to-noise ratio by systematically adding covariate-specific white noise to the input data. Each experiment was repeated five times for robust performance estimation and corrected for confounding age and sex effects.

**Figure 1:**
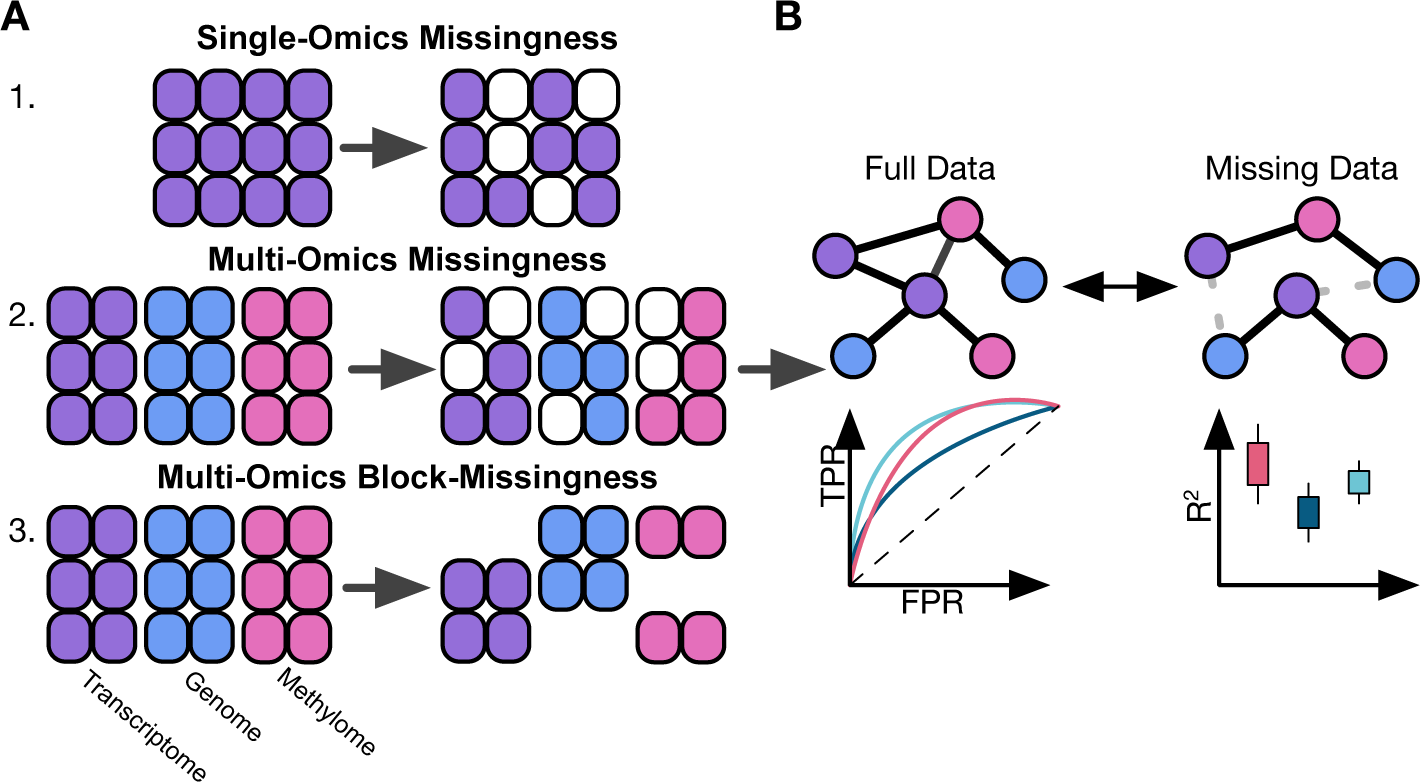
Benchmark schematic. **A)** Three common missingness scenarios in regulatory network inferences are tested: 1) single-omics, 2) multi-omics random missingness of individual elements, and 3) block-wise missingness in which entire omics layers are missing for an individual. **B)** We tested five approaches of two categories: 1) Two-step approaches that first impute and then aggregate imputation through, and 2) Inverse covariance estimation approaches that implicitly handle missingness during inference. We inferred regulatory networks from full data and data missing for each method and compared the resulting networks with multiple performance metrics.

#### Single-Omics missing

We selected the transcriptomics level as a single-omics layer to test the different models’ capabilities to infer gene regulatory networks with less directly informative co-correlation structures that could be used to impute the missing gene expression information. We removed *m* ∈ {*0%, 10%, 20%, 30%, 40%, 50%*} randomly selected entries from the input data. Additionally, we added noise with increasing intensity to the data by drawing from a normal distribution 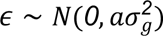 per gene with a gene-specific variance term estimated from the real data and *a* ∈ {*0, 0.5, 1.5*}.

#### Multi-Omics missing

To test the models’ capabilities to handle more complex co-correlation structures that could potentially be exploited for better imputation, we expanded the single-omics experiment to jointly consider the three available omics types. As before, we randomly removed *m* ∈ {*0%, 10%, 20%, 30%, 40%, 50%*} of entries independently per omics layer and added feature-dependent noise to the data as described before while ensuring to bound the beta values of the methylation data to the range between 0 and 1.

#### Multi-Omics Block-Missing

To test the capabilities of the method to handle block-wise missing information, we removed *m* ∈ {*0%, 10%, 20%, 30%, 40%, 50%*} samples per omics layer such that at least two omics-layers still remained per sample. Additionally, we added noise to the remaining samples as described before.

#### Downsampling

Similarly, to identify the minimal number of samples required to infer reliably the regulatory network, we downsampled the dataset to *k* ∈ {*90%, 80%, 70%, 60%, 50%*} of samples.

#### Runtime

Using TCGA-BRCA dataset, we tested the runtime of each method on a dedicated machine with an AMD EPYC 7502P 32-Core Processor with 2.5GHz base clock speed and 860 GB RAM using the multi-omics missingness experiment with the same configurations as before. We ran all experiments with ncores=60 (with hyperthreading enabled).

### 2.4 Evaluation Metrics

To construct the final network from the individual regressions, we applied a strict filter connecting independent to dependent variables if their beta coefficient was non-zero and their R^2^ > 0.1.

#### Prediction Metrics

To measure the prediction qualities of each regression model, we recorded the root means squared error (RMSE) and R^2^ respectively, and compared their distributions based on a Wilcoxon rank-sum test.

#### Network Reconstruction Metrics

To measure the methods’ abilities to handle missing data well, we compared the inferred regulatory networks from missing data to their counterpart inferred on full data, and calculated precision, recall, and F1-scores of the recovered network edges.

Similarly, we inferred a ground truth network with KiMONo using stability selection by repeating the network inference 100 times fitting different random seeds and averaging over the resulting coefficients and R^2^ values before constructing the network, improving the robustness of the graph resulting in a final network consisting of 2,458 nodes (2,182 genes, 63 CNVs, 211 methylations) and 6,554 edges for TCGA-BRCA dataset and 33,652 nodes (12,738 genes, 11,052 CNVs, and 9,862 methylations) and 87,229 edges for TCGA-MIBC dataset. We compared each stability-selected reference network to all inferred networks on missing data to determine performance differences across the individual methods.

#### Topological Metrics

The interpretation of complex heterogeneous networks and identification of key modules and important network nodes relies on the topological network features. Hence, it is also vital to evaluate if the methods can robustly infer topological structures. Therefore, we use multiple network-based metrics such as node-degree distribution, betweenness centrality, and clustering coefficient to quantify and compare the topological changes of the networks inferred from missing data. Node degree indicates the sparseness of the network, while betweenness centrality indicates how interconnected the network is, and the global clustering coefficient (transitivity) indicates how densely connected neighbouring nodes are.

#### Quantitative trait analysis

The sparse methods implicitly computed multivariate expression quantitative trait methylation (eQTM) as we mostly restricted the calculation between gene expression and their corresponding methylation site measurements. Hence, we compared our findings with a state-of-the-art method based on a linear regression approach to eQTM calculation, Matrix eQTL [49]. Our results enabled us to retrieved the genes connected to methylation sites based on topologies constructed using previously chosen thresholding: i) R-squared per model > 0.1, and ii) regression coefficient different to zero. For Matrix eQTL, we retrieved the significantly linked genes to eQTMs detected running both, imputed gene expression and imputed methylation sites measurements as inputs for the set of features in the different inferred networks, with a significance threshold per model of 1e-5, and FDR threshold of 0.01.

## Results

### Most topological features can be conserved in data with missingness

To investigate to what extent topologies of inferred networks are affected by noise and missingness, we computed multiple network properties across all benchmarking scenarios for all methods.

In single-omics scenarios, knnSGLasso demonstrated the most stable topology with increasing missingness rate on TCGA-BRCA data (Figure 2A&B, Supplementary Figures 2 & 3), decomposing into small subnetwork at high missingness rates according to the increase in transitivity (0.054±0.007, missingness = 0.1; 0.067±0.003, missingness = 0.5) and reduction in betweenness centrality (0.005±0.007, missingness = 0.1; 0.002±0.004, missingness = 0.5; Figure 2A&B). All methods were topologically more stable on TCGA-MIBC data except for CoCoLasso and HMLasso whose transitivity and betweenness centrality dropped at high missingness (transitivity: 0.077±0.002, betweenness: 0.003±0.002, missingness = 0.4; transitivity: 0.059±0.004, betweenness: 0.002±0.002, missingness = 0.5 for CoCoLasso; Figure 2A&C, Supplementary Figure 4). TCGA-PRAD showed similar topological results to TCGA-MIBC including the dropping of CoCoLasso and HMLasso at a high missingness rate (transitivity: 0.066±0.005, betweenness: 0.006±0.003, missingness = 0.5 for CoCoLasso, Supplementary Figure 8).

**Figure 2:**
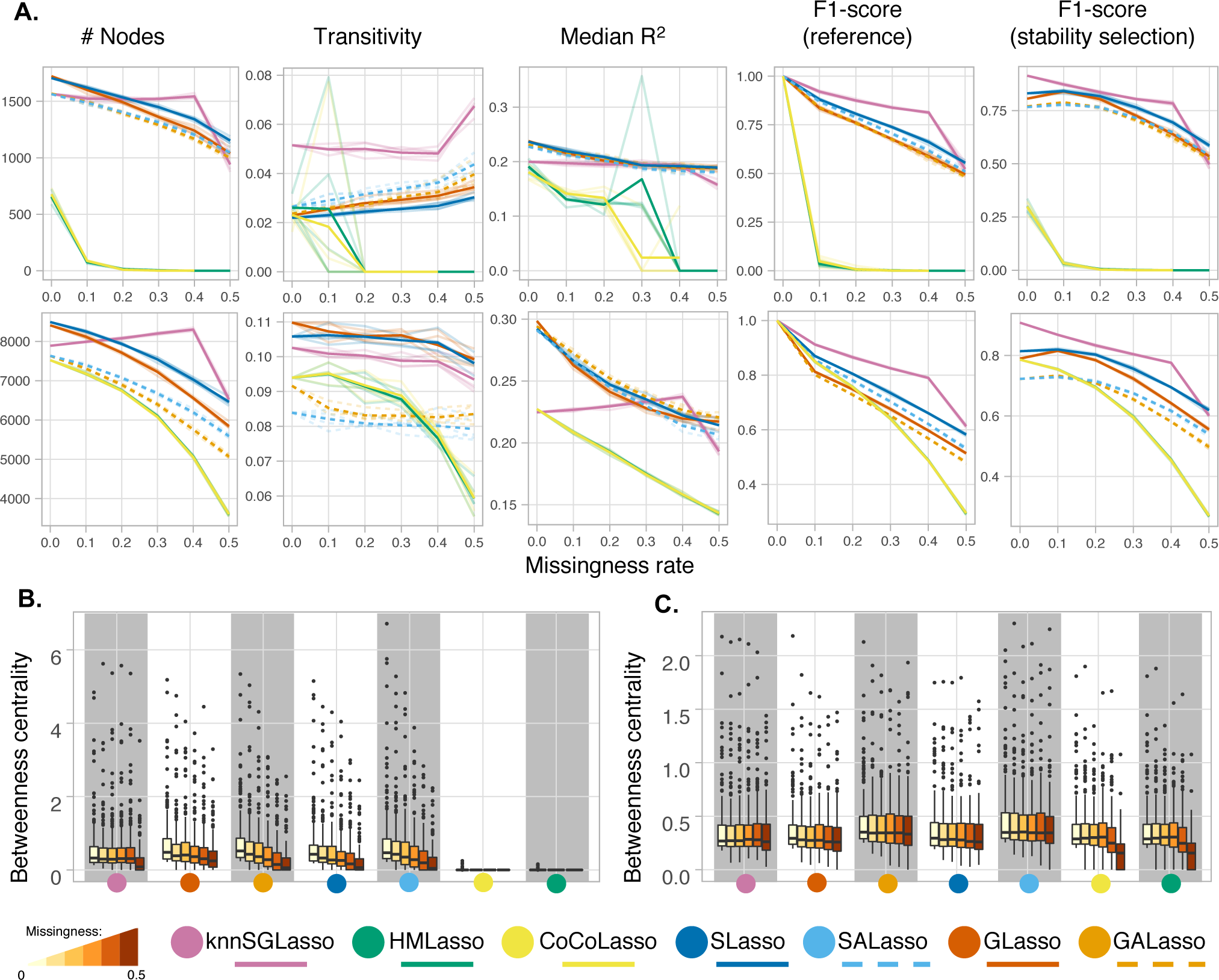
Benchmark results across single-omics experimental setups for both datasets. Transparent lines denote individual runs, while bold lines refer to the average performance. **A)** TCGA-BRCA dataset (top), TCGA-MIBC dataset (bottom). Performance was evaluated using network size (number of nodes), transitivity (global clustering coefficient), median R^2^, and F1-scores compared to a reference (i.e., the same method applied on the full data and stability selected networks generated with KiMONo). **B)** and **C)** Illustrating the topological change in TCGA-BRCA **B)** and TCGA-MIBC **C)** datasets scaling values for the top 200 nodes with the highest betweenness centrality through increased missingness.

In multi-omics scenarios, knnSGLasso had the most stable topology across increasing missingness rates on TCGA-BRCA. However, in presence of the highest missingness rate, knnSGLasso suffered an abrupt decay in betweenness centrality (Figures 3A&B). In contrast, knnSGLasso networks became sparser as indicated by the increase in the betweenness centrality and the slight reduction of transitivity with increasing missingness on TCGA-MIBC and TCGA-PRAD data (Figure 3A&C, Supplementary Figures 3, 8 & 9). S(A)Lasso and G(A)Lasso inferred networks on TCGA-MIBC and TCGA-PRAD data decreased their betweenness and transitivity abruptly compared to networks inferred on TCGA-BRCA data (Figures 3B&C; Supplementary Figures 3, 8 & 9). CoCoLasso and HMLasso slightly decreased their betweenness centrality for no to medium missingness rates (CoCoLasso and HMLasso: 0.002±0.001, for missingness = 0 / 0.3), and dropped in the presence of high missingness (CoCoLasso and HMLasso: 0.001±0.001, missingness = 0.5; Figure 3C; Supplementary Figure 9).

**Figure 3:**
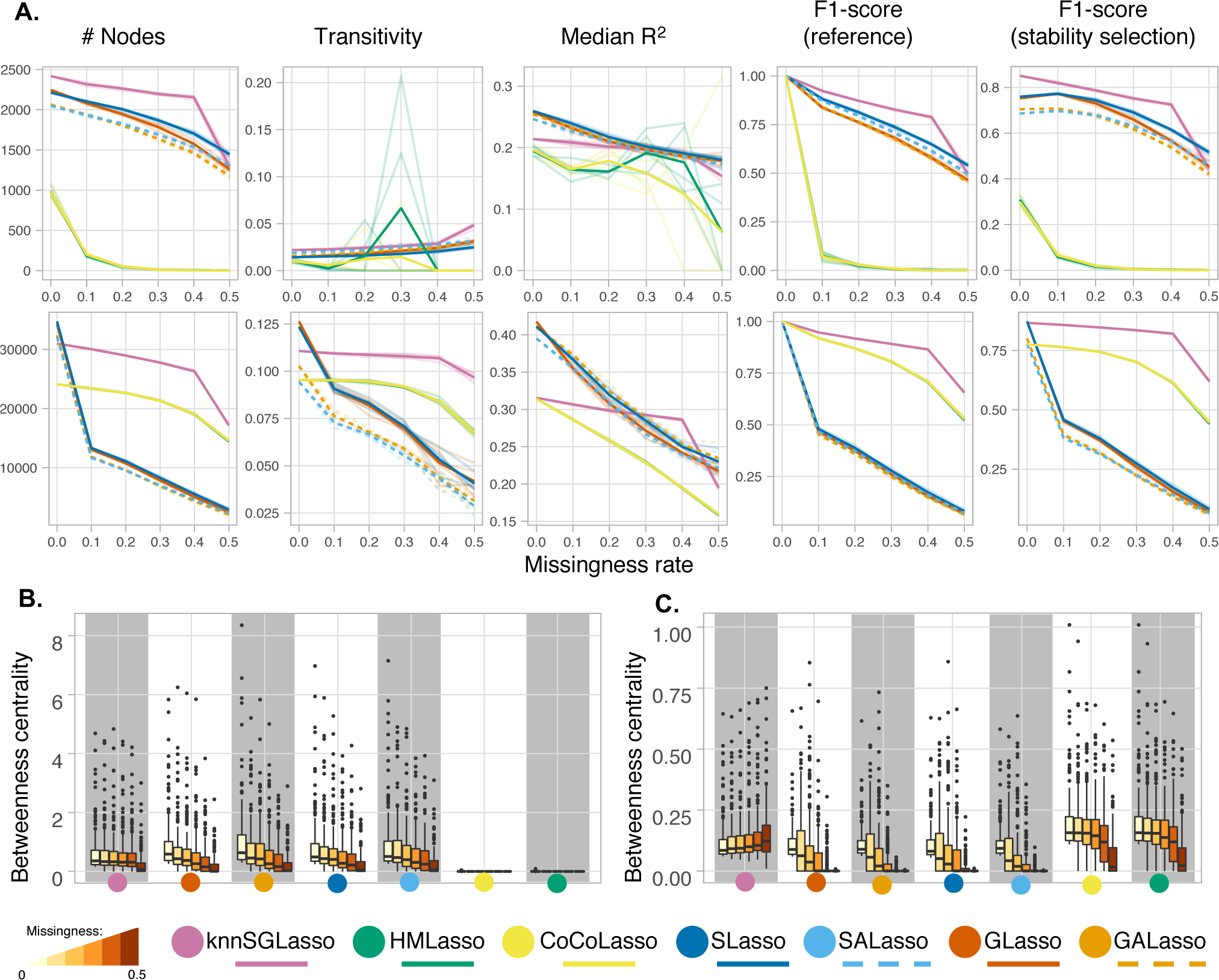
Benchmark results across multi-omics experimental setups for both datasets. Transparent lines denote individual runs, while bold lines refer to the average performance. **A)** TCGA-BRCA dataset (top), TCGA-MIBC dataset (bottom). Performance was evaluated using network size (number of nodes), transitivity (global clustering coefficient), median R^2^, and F1-scores compared to a reference (i.e., the same method applied on the full data and stability selected networks generated with KiMONo). **B)** and **C)** Illustrating the topological change in TCGA-BRCA **B)** and TCGA-MIBC **C)** datasets scaling values for the top 200 nodes with the highest betweenness centrality through increased missingness.

knnSGLasso and SALasso presented the most stable topologies in presence of block-missingness on TCGA-BRCA dataset (Figure 4B), although knnSGLasso’s full-data topology varied considerably compared to the counterpart networks inferred in presence of missing data (betweenness: 0.006±0.007 and 0.005±0.007, transitivity: 0.022±0.0 and 0.050±0.006, for missingness=0 and missingness=0.5, respectively; Figures 4A&B; Supplementary Figure 3). S(A)Lasso and G(A)Lasso were less stable, declining slightly in betweenness centrality with increasing missingness rate (Figure 4B). knnSGLasso behaved similarly on TCGA-MIBC and TCGA-PRAD data, the betweenness centrality (0.001±0.001, in both) was drastically different than the rest of its own inferred topologies in presence of missing data (0.004±0.003 and 0.004±0.004 for missingness = 0.1, respectively). The betweenness centrality for the approaches dealing with multiple imputed data dropped abruptly near to zero even at 10% of missingness (Figure 4C; Supplementary Figure 9). CoCoLasso and HMLasso had the most stable topologies including the full-data inferred network. However, their betweenness and transitivity were lower than knnSGLasso (CoCoLasso and HMLasso: 0.002±0.001 and 0.096±5e-4, knnSGLasso: 0.003±0.003 and 0.099±0.001, respectively for missingness=0.5; Figure 4A&C; Supplementary Figures 3 & 9).

**Figure 4:**
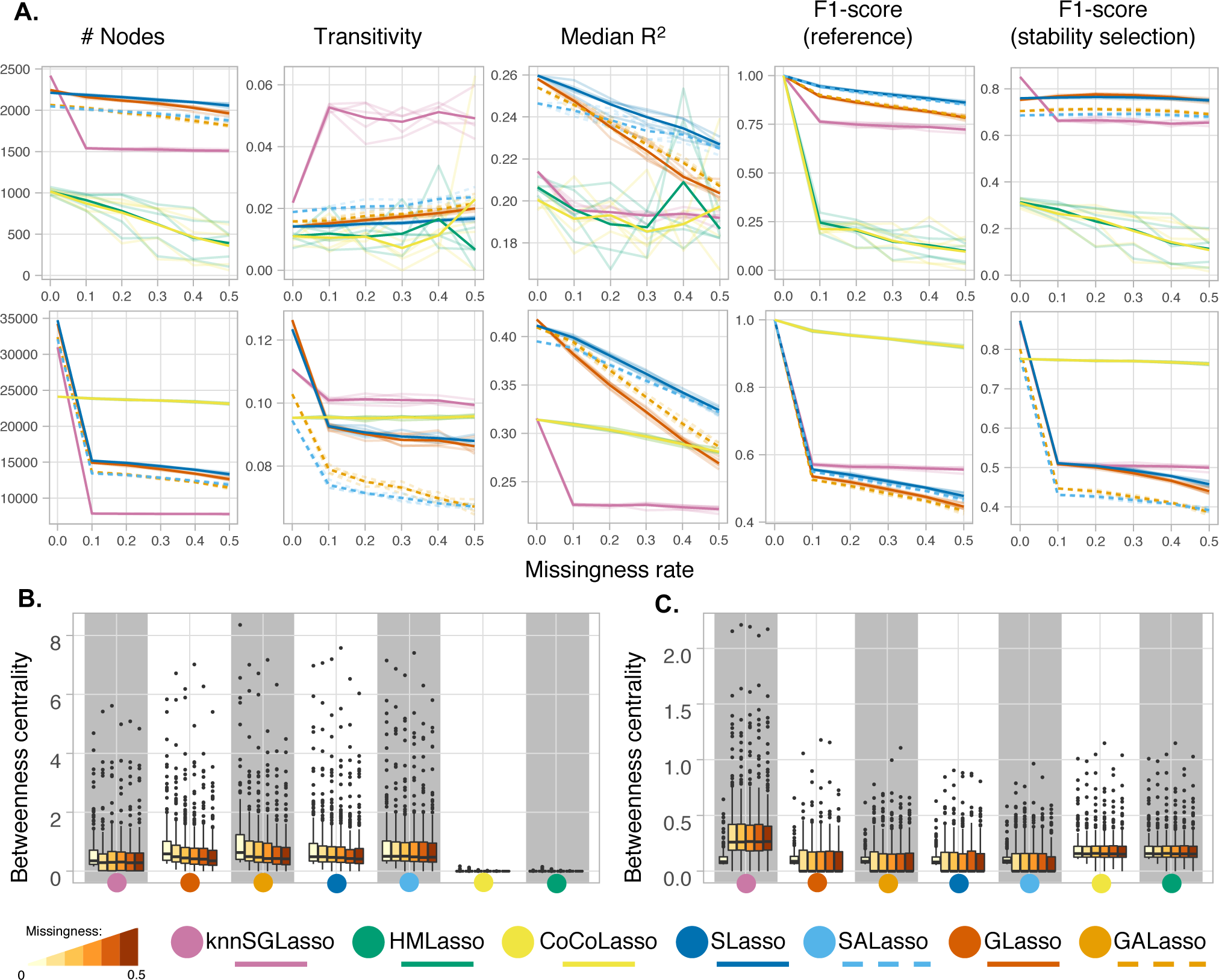
Benchmark results across block-missingness experimental setups for both datasets. Transparent lines denote individual runs, while bold lines refer to the average performance. **A)** TCGA-BRCA dataset (top), TCGA-MIBC dataset (bottom). Performance was evaluated using network size (number of nodes), transitivity (global clustering coefficient), median R^2^, and F1-scores compared to a reference (i.e., the same method applied on the full data and stability selected networks generated with KiMONo). **B)** and **C)** Illustrating the topological change in TCGA-BRCA **B)** and TCGA-MIBC **C)** datasets scaling values for the top 200 nodes with the highest betweenness centrality through increased missingness.

Across all benchmark settings, the methods showed a decreasing network size with increasing missingness. The constant loss of edges showed a stable ratio indicating no bias towards a specific omics layer (Supplementary Figures 4 & 8). GLasso was less affected in terms of noise, while knnSGLasso produced the most robust results in terms of missingness (Supplementary Figure 2).

### kNN imputation-based model performs best for single- and multi-omics data with random missingness

In both the single- and multi-omics setting, knnSGLasso was the best performing method independently of the dataset, reaching F1-scores of 0.921±0.005 and 0.912±0.001 on the single omics and 0.923±0.002 and 0.947±0.001 on the multi-omics data at 10% missingness (TCGA-BRCA and TCGA-MIBC data, respectively). Followed by SLasso (single-omics: 0.880±0.005 and 0.870±0.002) for single-omics, and CoCoLasso (0.919±0.001) for multi-omics when compared to the reference networks inferred on full data (Figures 2A & 3A; Supplementary Figures 5 & 6).

The performance gradually decreased with increasing missingness and noise levels to a similar degree for all the methods except for the specific case of multiple imputation-based methods in the multi-omics experiment on TCGA-MIBC data. In this particular case, the F1-score increased to 0.485±0.003 for SLasso, 0.442±0.004 for SALasso, 0.405±0.005 for GLasso, and 0.398±0.004 for GALasso, at 50% missingness and a medium noise (a=0.5; Supplementary Figure 6). At higher noise levels, the performance of all methods declined dramatically reaching only F1-scores below 0.15.

For the TCGA-BRCA dataset, HMLasso and CoCoLasso came in last, reaching F1-scores below 0.1 both on single- and multi-omics data even for samples with only 10% missingness and no noise (Figure 2A; Supplementary Figure 6). Both methods even failed to infer networks in 11/15 and 11/15 single-omics experiments at 30% and 40% missingness, as well as in 9/11 and 8/11 multi-omics experiments, at 40% and 50% missingness, respectively.

Similar behaviour could be observed when comparing the inferred networks to the stability selection-based reference network: knnSGLasso performed best with an F1-score of 0.873±0.005 and 0.869±0.008 on single-omics, and 0.820±0.004 and 0.858±0.007 on multi-omics at 10% missingness and no noise in the TCGA-BRCA and TCGA-MIBC dataset, respectively, followed by SLasso (TCGA-BRCA: 0.841±0.007, TCGA-MIBC: 0.820±0.003), for single-omics, and SLasso (TCGA-BRCA: 0.772±0.003) and CoCoLasso (TCGA-MIBC: 0.764±0.001) for multi-omics random missingness experiment (Figure 2A & 3A; Supplementary Figure 7). SLasso did marginally outperform knnSGLasso in the TCGA-BRCA dataset in scenarios with high noise and/or high missingness and was almost equivalent at high noise and high missingness (50% missingness, a=0.5) in the multi-omics scenario on the TCGA-MIBC dataset. At the highest noise levels (a=1.5), even in the absence of missingness, none of the methods exceeded F1-scores of 0.18 (Supplementary Figure 7).

TCGA-PRAD showed similar results to TCGA-MIBC in both single- and multi-omics (Supplementary Figure 10). At the single-omics level, knnSGLasso outperformed the rest of the methods in both low- and high-missingness based on F1-scores calculated using full data as reference (0.937±0.001 and 0.912±0.001 at 10% missingness; 0.71±0.017 and 0.613±0.004 at 50% missingness, respectively). Whilst CoCoLasso and HMLasso persisted as worst performing methods in presence of high missing ratio (0.488±0.004 and 0.492±0.004 for TCGA-PRAD, respectively; 0.293±0.004 and 0.295±0.004 for TCGA-MIBC, at 50% missingness) (Supplementary Figure 10). On the other hand, F1-score calculated based on a stable network at a high missingness ratio showed a slight difference between both datasets, in TCGA-MIBC, SLasso slightly outperformed knnSGLasso (0.619±0.007 and 0.601±0.003 at 50% missingness, respectively). TCGA-PRAD showed the opposite result (0.662±0.002 and 0.702±0.018 at 50% missingness, respectively) (Supplementary Figure 10).

At multi-omics level, knnSGLasso outperformed the rest of the methods followed by CoCoLasso and HMLasso in both, TCGA-MIBC and TCGA-PRAD in low- and high-missingness ratio according with F1-score calculated using whole data as reference (0.947±0.001, 0.919±0.001, 0.919±0.001, respectively at 10% missingness, and 0.656±0.001, 0.524±0.006, 0.528±0.005, respectively at 50% missingness for TCGA-MIBC; 0.955±0.001, 0.930±0.001, 0.930±0.001, respectively at 10% missingness, and 0.697±0.017, 0.593±0.006, 0.596±0.005, respectively at 50% missingness for TCGA-PRAD), and the same measurement using stable network as reference instead (0.858±0.001, 0.764±0.001, 0.764±0.001, respectively at 10% missingness, and 0.620±0.002, 0.443±0.005, 0.447±0.005, respectively at 50% missingness for TCGA-MIBC; 0.874±0.001, 0.778±0.001, 0.778±0.001, respectively at 10% missingness, and 0.663±0.016, 0.506±0.005, 0.509±0.005, respectively at 50% missingness for TCGA-PRAD) (Supplementary Figure 10).

Adaptive weights had a marginal impact on performance with a slight negative effect, irrespective of the dataset. Only at high noise levels did adaptive weights stabilise performance (Supplementary Figures 6, 7 & 10).

### SLasso and inverse covariance-based approaches perform best for data with block-missingness

When investigating block-missingness, i.e., where entire omics layers are missing for a subset of samples, SLasso performed best when TCGA-BRCA data were used, while CoCoLasso and HMLasso performed best on TCGA-MIBC and TCGA-PRAD data (Figure 4; Supplementary Figures 2 & 10). SLasso, CoCoLasso, and HMLasso reached high consistency with the networks inferred on full data with SLasso achieving F1-scores between 0.945±0.004 at 10% missingness (a=0) and 0.688±0.002 at 50% missingness (a=0.5) on TCGA-BRCA data. CoCoLasso and HMLasso reached F1-scores of 0.967±0.002 at 10% missingness (a=0), and 0.810±0.004 at 50% missingness (a=0.5) respectively on TCGA-MIBC input data (Figure 4A; Supplementary Figure 5 & 6). The results obtained for TCGA-MIBC were consistently similar to those obtained for TCGA-PRAD with F1-scores of 0.976±0.002 at 10% missingness (a=0), and 0.920±0.004 for TCGA-MIBC and 0.938±0.005 for TCGA-PRAD at 50% missingness (a=0) (Supplementary Figure 10).

With higher noise, the performance dropped below an F1-score of 0.2 (Supplementary Figure 6). This behaviour could also be observed for GALasso and knnSGLasso, although with generally lower F1-scores. Notably, knnSGLasso networks almost exclusively consisted of gene nodes, while all other methods had a proportional representation of all omics types. The implementation of kNN-imputation used here is not able to handle entire block-missing samples and, consequently, knnSGLasso removes a substantial amount of the features.

Similar behaviour could be observed when comparing the inferred networks on missing data to the stability-selection-based reference. Both SLasso and GLasso outperformed all other methods on TCGA-BRCA data with SLasso reaching the highest F1-scores of 0.763±0.003 (10% missingness, a=0) to 0.655±0.007 (50% missingness, a=0.5), while CoCoLasso and HMLasso outperformed all methods on TCGA-MIBC data reaching F1-scores of 0.773±0.001 (10% missingness, a=0) to 0.728±0.003 (50% missingness, a=0.5; Figure 4A). This same pattern was observed on TCGA-PRAD with F1-scores of 0.788±0.001 at 10% missingness (a=0), and 0.763±0.002 for TCGA-MIBC and 0.778±0.002 for TCGA-PRAD at 50% missingness (a=0) (Supplementary Figure 10). Adaptive weights (SALasso and GALasso) again had negligible effects and only improved performance markedly at high noise levels as observed in TCGA-MIBC (a=1.5; Supplementary Figure 7).

HMLasso and CoCoLasso performed worst on TCGA-BRCA data, reaching average F1-scores of 0.212±0.022, and 0.246±0.039 at 10% block-missingness, dropping to 0.063±0.033 and 0.073±0.031 respectively with 50% block-missingness and medium noise (a=0.5; Supplementary Figure 6). Comparing those methods to the stability-selection-based reference depicted a similar picture (Supplementary Figure 7).

### knnSGLasso is the least affected by sample size reduction

All methods showed a decrease in concordance already at 10% sample reduction (Figure 5). SLasso and knnSGLasso were the most stable method at single-omics level over progressive sample reduction (Figure 5). The F1-score of knnSGLasso was similar to SLasso on TCGA-BRCA data at 10% sample reduction (knnSGLasso: 0.906±0.018, SLasso: 0.906±0.003). However, at a high sample removal rate (50% sample reduction), SLasso (0.810±0.012) slightly outperformed knnSGLasso (0.796±0.020; Figure 5A). CoCoLasso and HMLasso reached similar performance as knnSGLasso on TCGA-BIMC and TCGA-PRAD data when 10% sample reduction (CoCoLasso and HMLasso: 0.916±0.005 and 0.949±0.005, knnSGLasso: 0.917±0.002 and 0.950±0.003, respectively), but were consistently outperformed by knnSGLasso at higher sample reduction rates (CoCoLasso and HMLasso: 0.782±0.005 and 0.849±0.005, knnSGLasso: 0.806±0.013 and 0.871±0.003, respectively; Figure 5A; Supplementary Figure 10). Similar behaviour but more prominently could be observed when comparing performance based on the stability-selection reference network irrespective of the dataset. knnSGLasso outperformed the rest of the methods followed by SLasso and GLasso. CoCoLasso and HMLasso were among the worst-performing methods (Figure 5A&B; Supplementary Figure 10).

**Figure 5.**
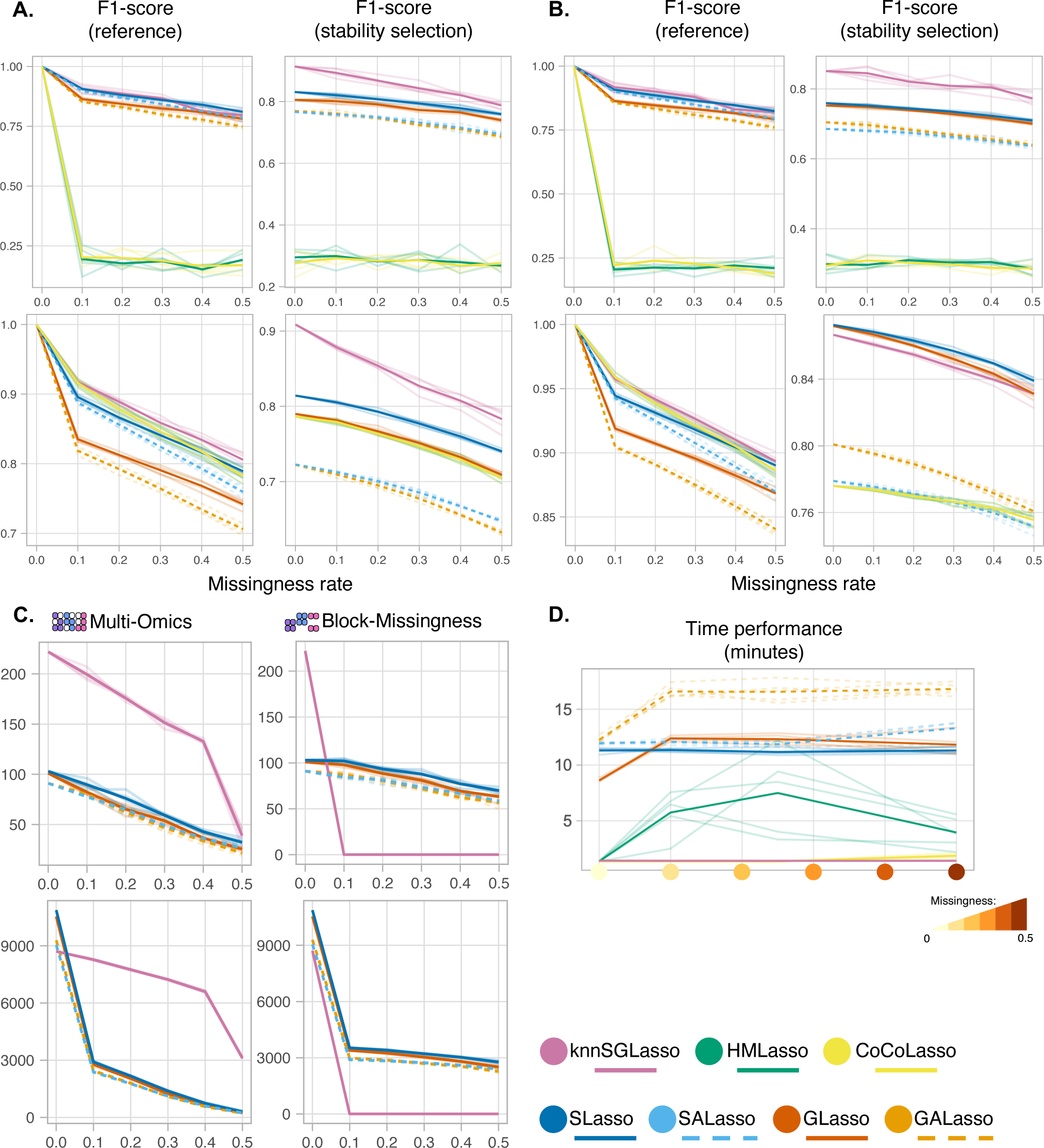
Sample size and inter-layer results across all experimental setups. Transparent lines denote individual runs, while bold lines refer to the average performance. F1-scores for single-omics **A)** and multi-omics **B)** on TCGA-BRCA data (top) and TCGA-MIBC data (bottom). **C)** eQTM-linked genes (number of genes with at least one methylation site significantly related) detected by Matrix eQTL based on the subset of nodes from sparse models inferred networks. **D)** Runtime evaluation over multi-omics experiments using TCGA-BRCA data.

On multi-omics data, the results showed a similar pattern: knnSGLasso consistently outperformed all other methods on TCGA-BRCA data (Figure 5B). On TCGA-MIBC and TCGA-PRAD data, CoCoLasso and HMLasso outperformed knnSGLasso at 10% sample reduction (CoCoLasso and HMLasso: 0.959±0.003 and 0.967±0.003, knnSGLasso: 0.958±0.001 and 0.966±0.002, respectively), but were overtaken by knnSGLasso with higher reduction rates (CoCoLasso and HMLasso: 0.886±0.005 and 0.907±0.005, knnSGLasso: 0.893±0.006 and 0.911±0.003, respectively; Figure 5B; Supplementary Figure 10). knnSGLasso strongly outperformed all other methods on TCGA-BRCA data when compared to the stability-selection-based reference network (Figure 5B). On TCGA-MIBC data, however, SLasso performed best, closely followed by GLasso. CoCoLasso and HMLasso performed worst for no to medium missingness rate, narrowly overtaking SALasso in presence of high missingness rate (CoCoLasso and HMLasso: 0.774±0.001 and 0.756±0.004; SALasso 0.776±0.001 and 0.752±0.004, for a=0.1 and a=0.5, respectively; Figure 5B). Compared to TCGA-MIBC, TCGA-PRAD had similar pattern results being SLasso best performing, closely followed by GLasso in low to high missingness rate (0.877±0.001 and 0.875±0.001 at 10% missingness, 0.854±0.001 and 0.848±0.003 at 50% missingness, respectively). However, at no-missingness, knnSGLasso narrowly overtook SLasso and GLasso (SLasso and GLasso: 0.879±0.0, knnSGLasso: 0.883±0.0). CoCoLasso and HMLasso performed worst in most missingness rates (Supplementary Figure 10).

### Inter-layer links possess biological meaning comparable to state-of-the-art methods

We have successfully identified inter-layer links representing potential biological interactions between different molecular abstraction layers. Here, we performed multivariate regression to detect links between gene expression and methylation site measurements, which are expression quantitative trait methylation (eQTMs). To evaluate the viability of the eQTMs detected by Lasso models dealing with missing data, we compared our results against a state-of-the-art method based on linear regression to eQTM detection, Matrix eQTL [49] (Figure 5C). Matrix eQTL detected more eQTMs than sparse model-based networks, however, the eQTM-linked genes (number of genes with at least one methylation site significantly related) detected by Matrix eQTL were quite similar but marginally less than those eQTMs-linked genes detected by sparse models across the different experimental setups performed in this benchmarking (Figure 5C, Supplementary Figure 11). knnSGLasso results detected more eQTM-linked genes at the presence of multi-omics missingness in both datasets at a low missingness rate (199.4±5.03 for TCGA-BRCA and 8275.6±29.99 for TCGA-MIBC, at 10% missingness using Matrix eQTL; 202±4.90 for TCGA-BRCA and 8286.4±30.96 for TCGA-MIBC at 10% missingness using sparse models), as well as at high missingness rate (39.75±8.10 for TCGA-BRCA and 3117.4±45.84 for TCGA-MIBC, at 50% missingness using Matrix eQTL; 41.4 ±8.41 for TCGA-BRCA and 3305.8±44.95 for TCGA-MIBC, at 50% missingness using sparse models. Except at whole data performance where SLasso outperformed knnSGLasso followed by GLasso for TCGA-MIBC dataset (10875±0 for SLasso and 10540±0 for GLasso using Matrix eQTL; 10881±0 for SLasso and 10547±0 for GLasso using sparse models) (Figure 5C, Supplementary Figure 11).

At the presence of block-missingness, Matrix eQTL performance based on SLasso results detected the most eQTM-linked genes at low missingness rate (102±2.92 for TCGA-BRCA and 3527.2±12.76 for TCGA-MIBC, at 10% missingness using Matrix eQTL; 111±2.24 for TCGA-BRCA and 3552.6±12.52 for TCGA-MIBC, at 10% missingness using sparse models), as well as at high missingness rate (69.8±3.03 for TCGA-BRCA and 2771±61.53 for TCGA-MIBC, at 50% missingness using Matrix eQTL; 83±1.58 for TCGA-BRCA and 2886.8±54.41 for TCGA-MIBC, at 50% missingness using sparse models) (Figure 5C, Supplementary Figure 11).

### HMLasso is the fastest approach in datasets with no to medium missingness

knnSGLasso, HMLasso, and CoCoLasso had a similar runtime of 82 - 85 sec on the TCGA-BRCA dataset without missing information and gradually increased in runtime with rising missingness (Figure 5D). HMLasso was the fastest approach with an average runtime of 81.843±0.375 sec at low to medium missingness levels. Only in scenarios with high missingness, HMLasso was outperformed by knnSGLasso, reaching an average runtime of 97.089±1.327 sec. knnSGLasso demonstrated a very consistent runtime across all missingness levels with an average runtime of 97.725±7.995 sec. CoCoLasso behaved similarly but was affected by the degree of missingness, reaching maximum average runtimes of 449.6±225.2 sec. GALasso and SALasso were the slowest, with average runtimes of 739±15.532 sec and 919.785±11.833 sec on the complete dataset, respectively, of which 241.191±2.975 sec were dedicated to imputation. Overall runtime gradually increased for both methods reaching an overall average runtime of 1148.871±245.077 sec and 1184.688±157.794, respectively, of which on average 507.674±58.126 sec was spent in imputation. For GALasso, adaptive weights calculations added another 275.363±54.180 sec on average to the overall runtime.

## Discussion

Due to economic or technical restrictions, missingness of individual values or block-missingness of entire omics layers in a subset of samples is typical for high-throughput multi-omics experiments, rendering multi-omics network inference challenging. In this study, we benchmarked novel regression approaches that can handle missing information across common missingness scenarios in single- and multi-omics experiments and integrated these approaches into KiMONo, a recent approach for network-guided multi-omics network inference.

We observed that the performance of the different methods was dependent on the individual feature size per omics layer. However, some general trends could be identified: kNN-imputation combined with the standard KiMONo approach performed best in most cases except when block-missingness was present. For this particular case, the number of features per omic layer in the dataset was a crucial factor, with SLasso and GLasso being suitable for TCGA-BRCA data, while CoCoLasso and HMLasso were better suited for TCGA-MIBC and TCGA-PRAD data. Both types of methods were also able to handle high degrees of block-missingness consistently well on TCGA-BRCA, TCGA-MIBC and TCGA-PRAD data, respectively, providing an advantage in real-world applications.

One reason why multiple-imputation-based methods provided good performances in presence of low-dimensional omics-layers such as CNVs in TCGA-BRCA with 84 features, might be due to their utility of the posterior predictive distribution of the missing data and harmonised feature selection over the feature estimates resulting in robust selections more so than point estimates of inverse covariance-based methods. While the original paper of SALasso and GALasso claimed that adaptive weights improved the general performance [28], our results showed a marginal reduction. This discrepancy could be due to the lower imputation quality of the network-based multiple-imputation approach (ngMICE) we applied, propagating the imputation uncertainty into network inference. Increasing the number of multiple imputations could improve the performance, however, with an increase in runtime. Multiple imputation in high-dimensional data is generally challenging since existing approaches do not scale to the number of covariates typically encountered in multi-omics studies. Here, dimensionality reduction methods for multiple imputation [50], latent factor models [51], or deep-learning-based approaches [52–54] might improve multiple imputation and therefore network inference quality. However, a benefit of such explicit approaches is their ability to use and adequately address prior imputed datasets often provided by larger consortia.

Implicit methods relying on inverse covariance matrix estimation performed poorly on TCGA-BRCA data but performed generally well on TCGA-MIBC and TCGA-PRAD data, even outperforming knnSGLasso and SLasso/SALasso in presence of block-missingness, indicating that large volumes of information per omic-layer are required to estimate the covariance structure properly with such methods.

Inter-layer links with biological meaning were captured by the different methods across different missingness rates as the number of genes linked to methylation sites analysis demonstrated when compared to state-of-the-art methods such as Matrix eQTL [49] which is dedicated to eQTM detection. Matrix eQTL detected more eQTMs than sparse models-based networks as expected given this method uses inference based on linear regression for all possible combinations of genes-methylation sites as background procedure [49]. However, the number of total genes significantly involved in at least one eQTM was marginally less than those detected by the Lasso network inferred network in presence of missing data. Therefore, we proved the biological value of inter-layer links detected across different network inferences and showed the power of the methods to detect new biological insights. Furthermore, knnSGLasso conserved at least twice as many eQTMs-linked genes as the rest of the methods including when no-missingness data were used as input in TCGA-BRCA. The absence of advanced methods for multiple imputation, which optimally encompassed high imputation quality and low computational cost were factors that affected SLasso/SALasso and GLasso/GALasso in this task. However, the approach used here (ngMICE) demonstrated to be suitable across increasing missingness rates at the point that SLasso and GLasso outperformed the rest of the methods in certain conditions, such as block missingness in the TCGA-BRCA dataset.

Comparing similar datasets, such as TCGA-MIBC and TCGA-PRAD, and analysing a comparable dataset, such as TCGA-BRCA, through the implementation of different prior networks allowed for the characterisation of the strengths and weaknesses of different sparse models in network inference. We expect these findings to hold in other datasets, such as colorectal cancer (CRC) from TCGA used by Welz et al. [55] for network inference using KiMONo based on additional omics layers evaluated in the present benchmarking such as proteomics and DNA mutation [55]. This includes the improvement of recent frameworks such as DiffBrainNet, which includes differential expression analysis and prior-knowledge network inference (KiMONo) to study stress response in mouse brain [56].

We note that a true gold standard for evaluating the performance of network inference methods is missing. In its absence, we rely on networks inferred from complete, unperturbed data that nevertheless are likely to contain both false positive and false negative interactions which may affect the results. Hence, our reference networks are not suited for evaluating methods following different principles for inference as GENIE3 [57], or other nonlinear approaches such as ARACNe and their derivatives [7,8]. Nevertheless, the methods tested here can be used as direct substitutes for other Lasso-based network inference frameworks such as wgLasso [13], pLasso [58] or PoLoBag [59] with similar expected behaviour as demonstrated here. Also, other linear methods such as WGCNA [5] or Petal [60] could benefit from two-step approaches and consistent feature selection across multiple imputed datasets, similar to G(A)Lasso. A way of such approaches to handle multiple imputed datasets could be through the usage of Rubin’s rule after Fisher’s Z-transformation, which allows to pool the estimated correlation coefficients between omics features from multiple imputed datasets in a consistent manner [61,62].

In summary, we found explicit methods to be more robust than methods implicitly handling missingness in most scenarios except on block-missingness. While most methods were tolerant to high levels of missingness, they were strongly affected by noise. While HMLasso was the fastest tested method, knnSGLasso showed the best trade-off between performance and runtime and is thus our recommended approach for handling missingness in KiMONo except for consistently high-dimensional omics-layers data with block-missingness where we recommend HMLasso instead. While we see room for further method improvements, particularly with respect to multiple imputations of high-dimensional single matrices and the robustness of inverse covariance methods, our results show that robust multi-omics network inference in the presence of missingness is feasible with KiMONo and thus allows users to leverage available multi-omics data to their fullest extent.

## Key Points

- We extended KiMONo, a multi-omics network inference approach that is using prior network information, to handle data with missingness by integrating advanced Lasso-based inference techniques.
- knnSGLasso outperformed all other methods in the majority of scenarios except in presence of block-missingness where the kNN-imputation step tends to fail. In such cases, we recommend using SLasso on imbalanced and HMLasso on balanced omics layers dimensional data.
- Inverse covariance matrix estimation-based approaches were considerably sensible to the dimensionality of the input data, requiring high-dimensional data per omics layer to reach high performance.

## Contributions

**JDH, MiL, MA, AG** implemented the different Lasso approaches. **JDH** implemented the benchmark framework**. JDH, BS** analysed the results**. BS, CO, MaL** conceived and supervised the study. **JDH, BS, CO, MaL** wrote the original manuscript. **MaL, CO, FJT, BS, MiL** reviewed and edited the manuscript. All authors read and approved the final manuscript.

