## Supplementary file for "Multi-Omics Regulatory Network Inference in the Presence of Missing Data"

### Supplementary Figures

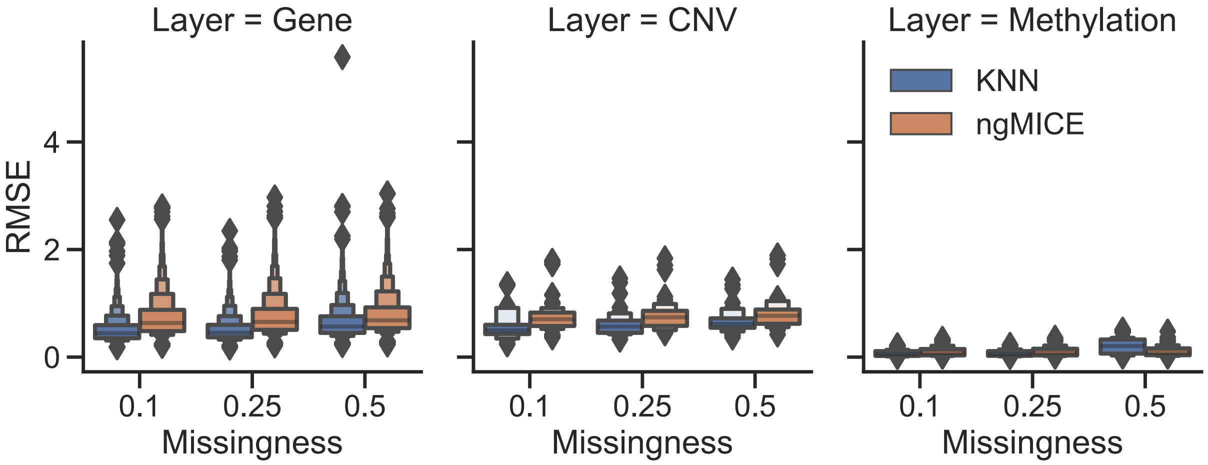


**Supplementary Figure 1.** Root means square mean error of kNN-based and network-based multiple imputation per omics layer with increased missingness (10%, 25%, 50%) removing randomly elements from each omics individually.

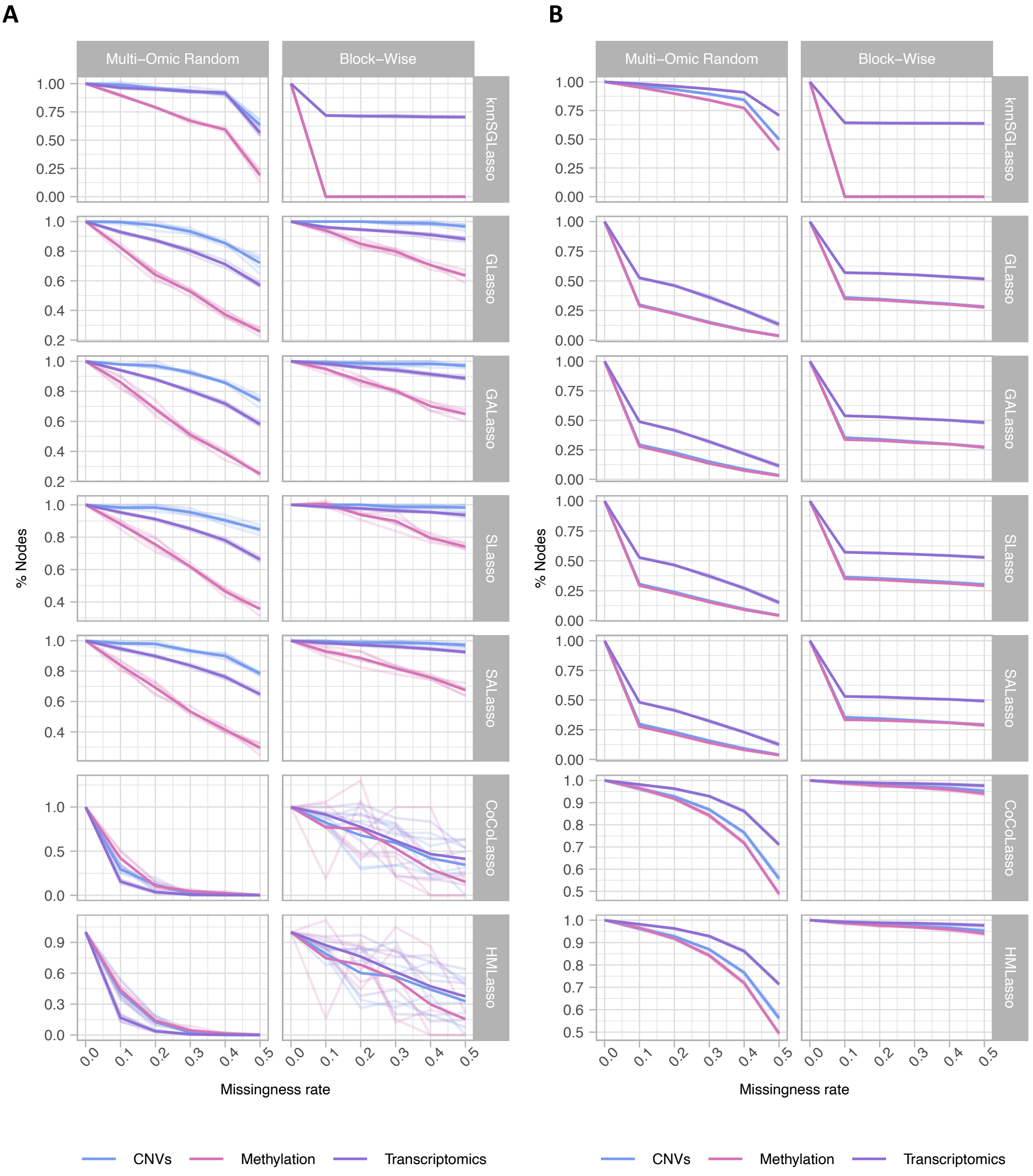


**Supplementary Figure 2.** Feature reduction per omics layer represented as the percentage loss compared to the full-data network inference, **A)** TCGA-BRCA, **B)** TCGA-MIBC.

**A**

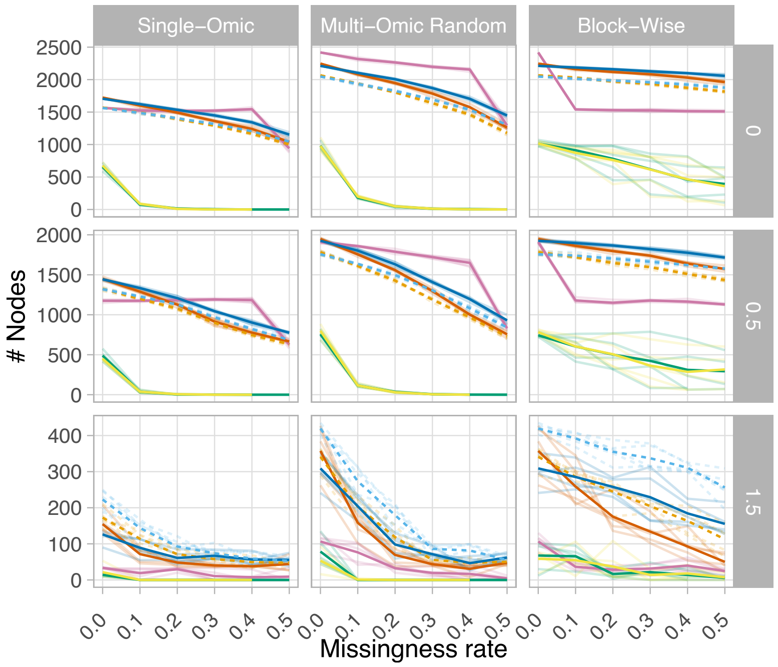


**B**

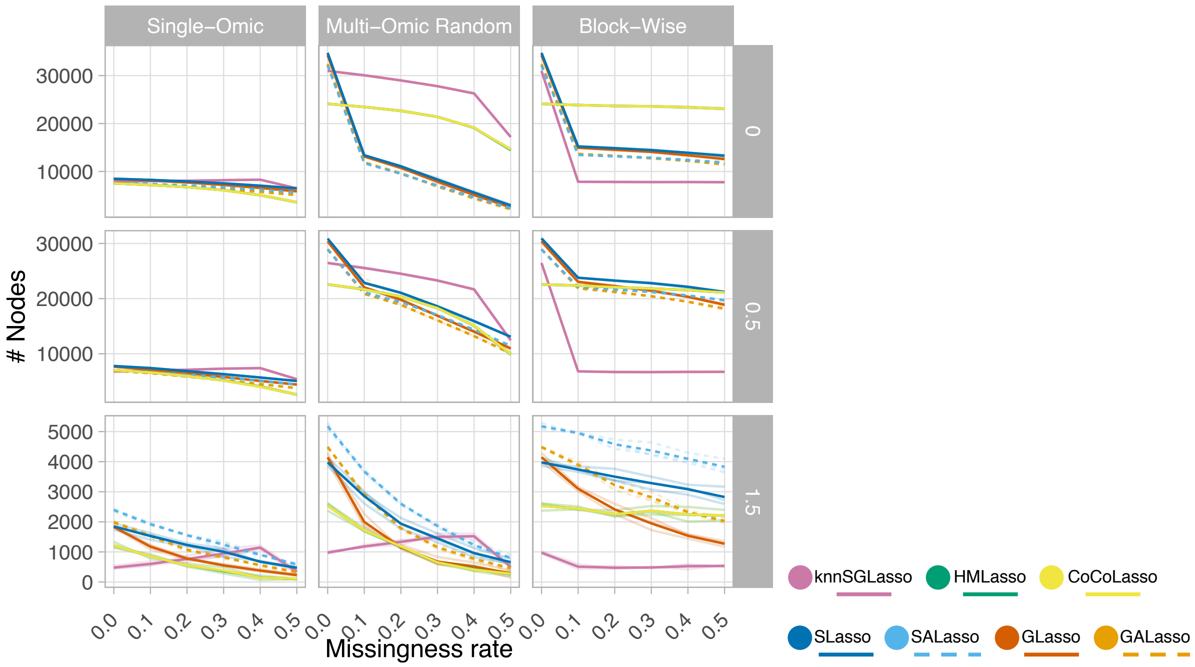


**Supplementary Figure 3.** Number of nodes indicating network size across all tested noise levels, **A)** TCGA-BRCA, **B)** TCGA-MIBC.

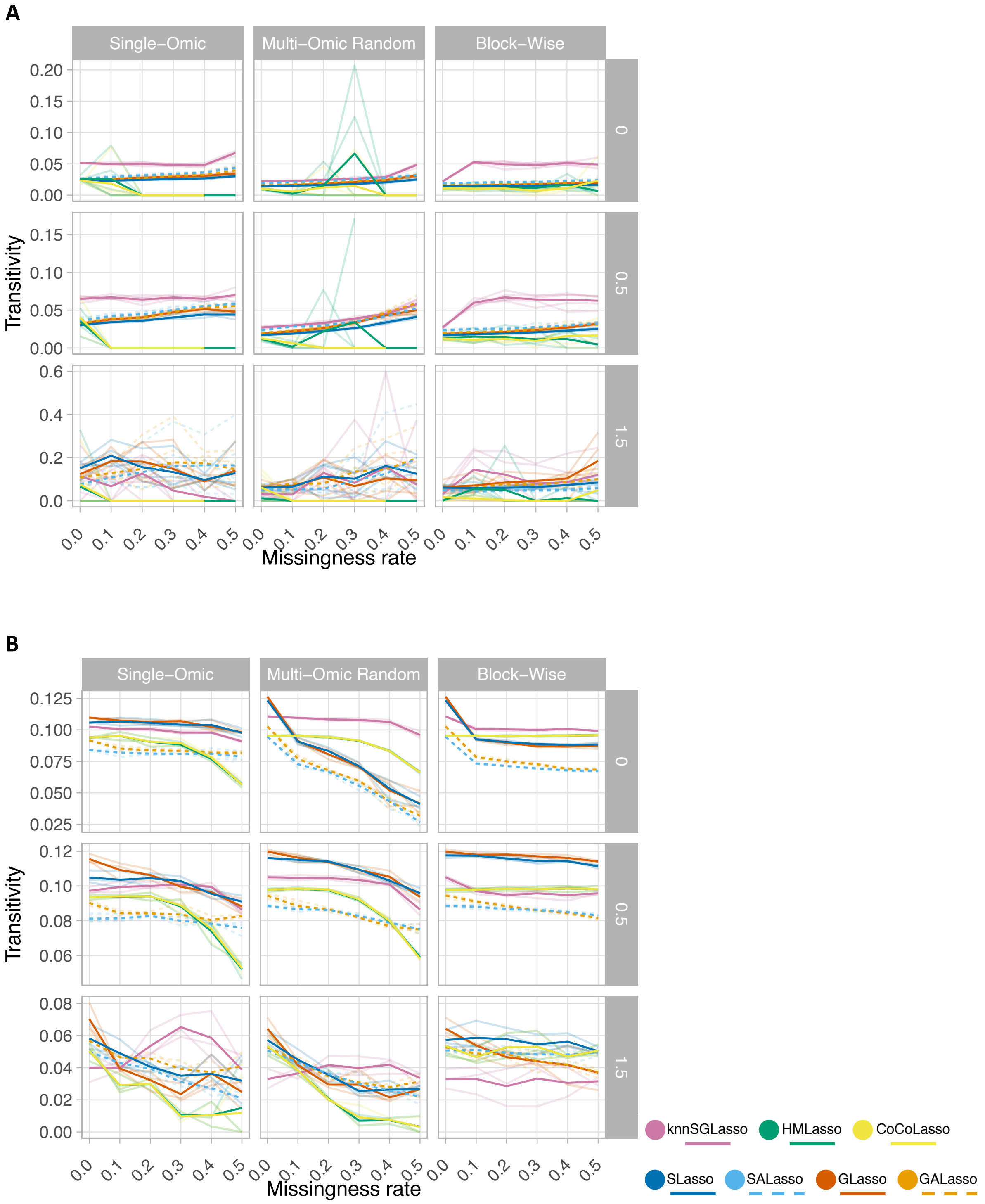


**Supplementary Figure 4.** Global clustering coefficient per network inferred. **A)** TCGA-BRCA, **B)** TCGA-MIBC.

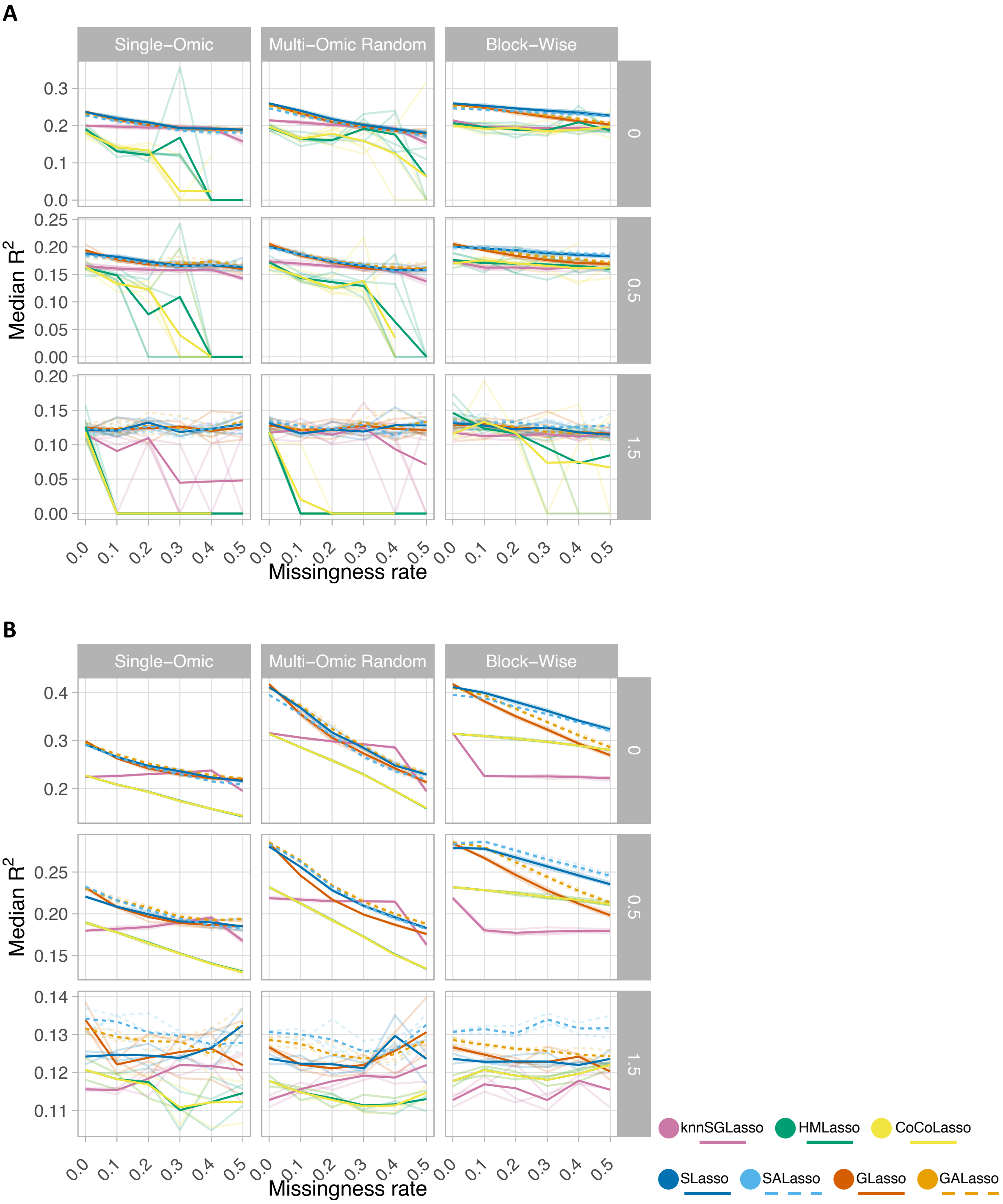


**Supplementary Figure 5.** Performance evaluation via $R^2$. For each network all nodes are evaluated via $R^2$. A single networks' quality can be estimated via median calculation across all models, **A)** TCGA-BRCA, **B)** TCGA-MIBC.

A

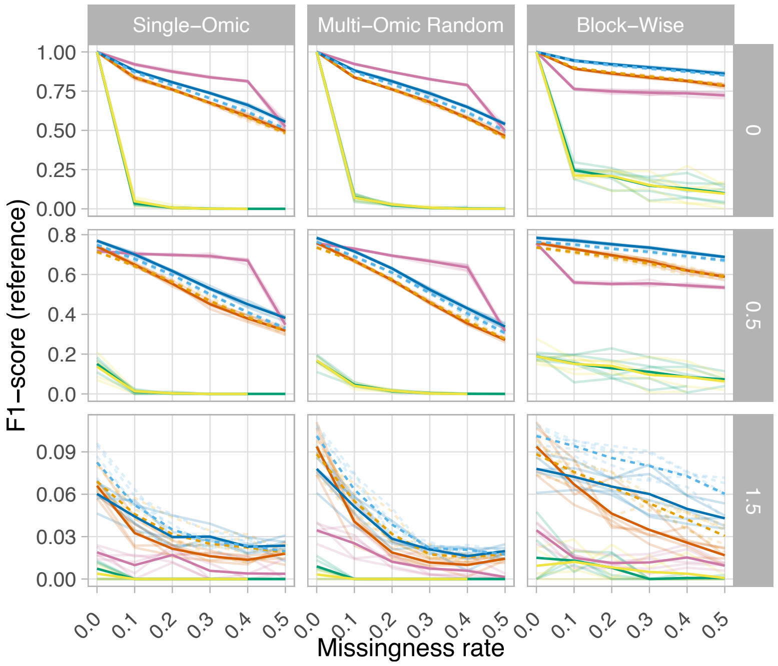


B

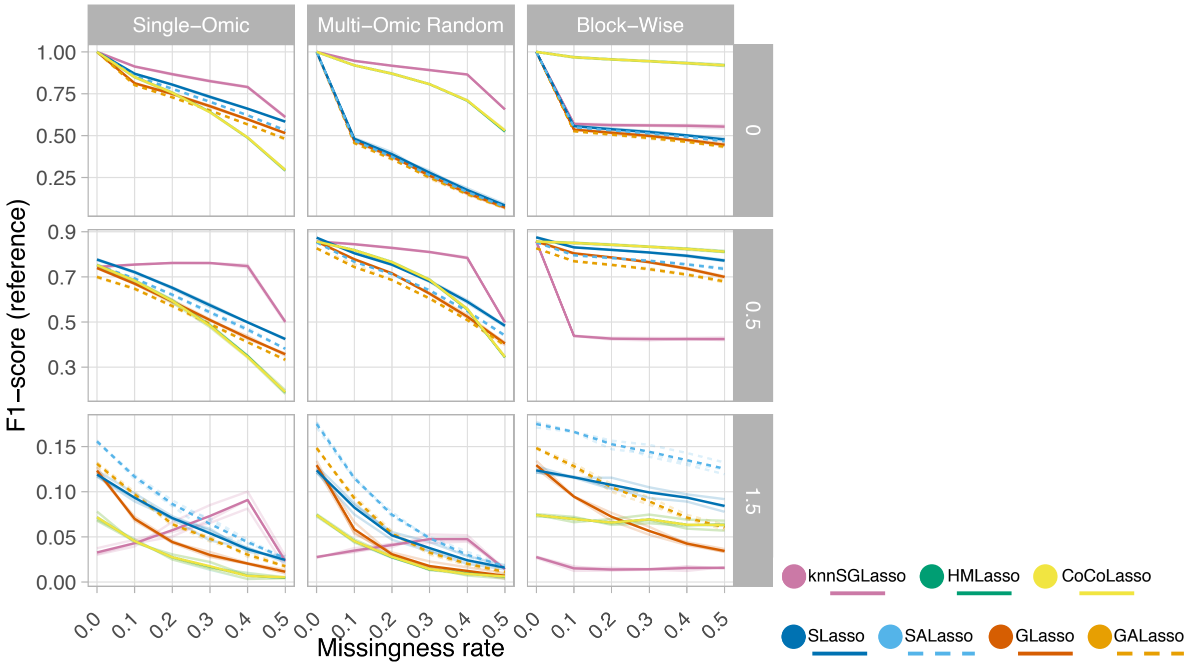


**Supplementary Figure 6.** Performance evaluation via F1-score. Network inference using full data was used as reference to calculate the F1-score per lasso model, **A)** TCGA-BRCA, **B)** TCGA-MIBC.

**A**

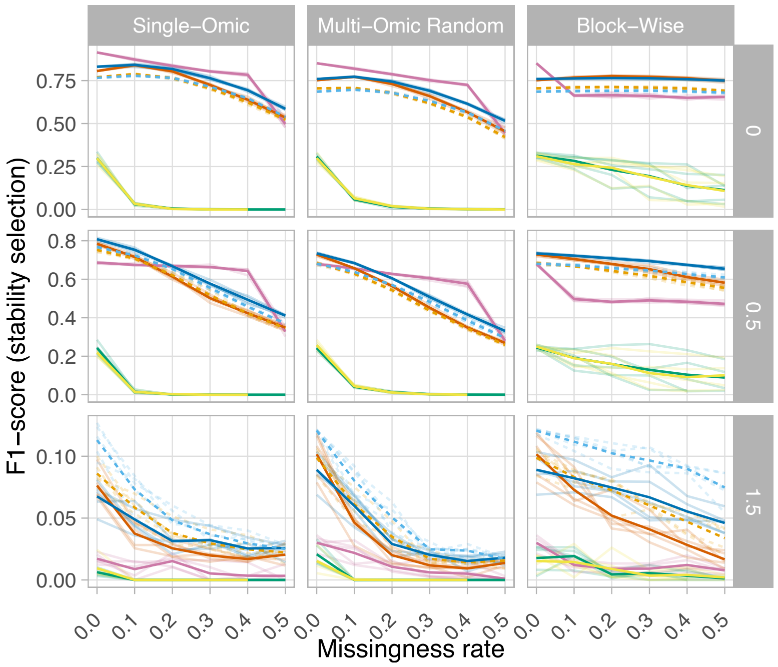


**B**

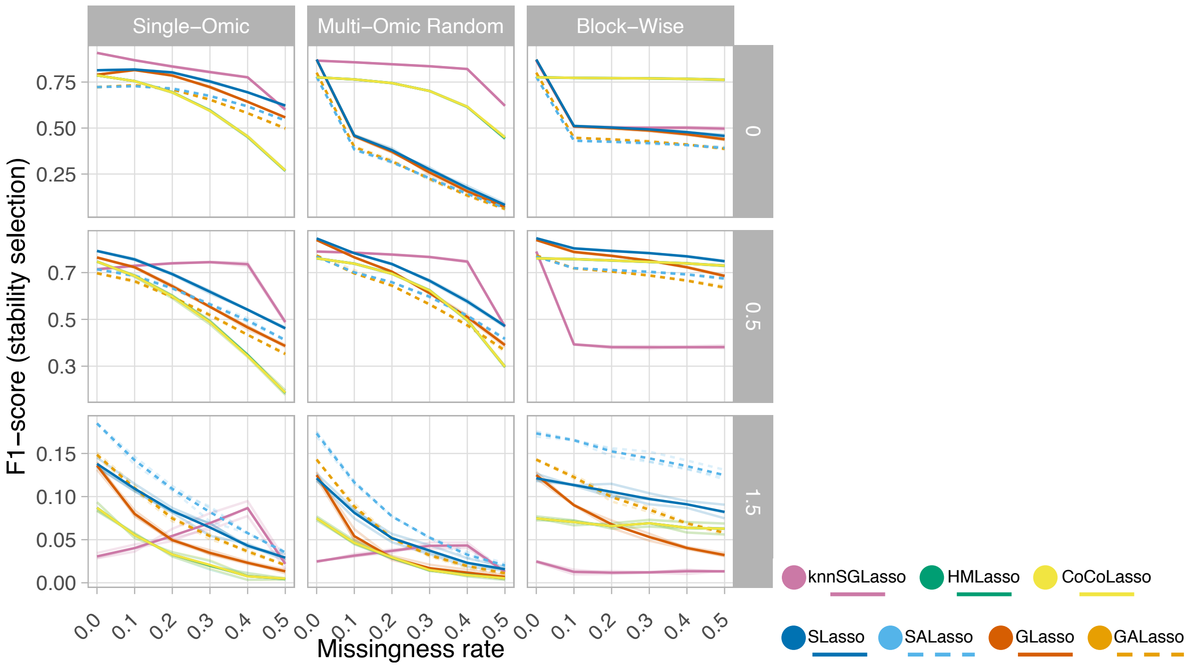


**Supplementary Figure 7.** Performance evaluation via F1-score. Network inference using KiMONo with stability selection (*k* = 100) over full data was used as reference to calculate F1-score for all lasso models, **A)** TCGA-BRCA, **B)** TCGA-MIBC.

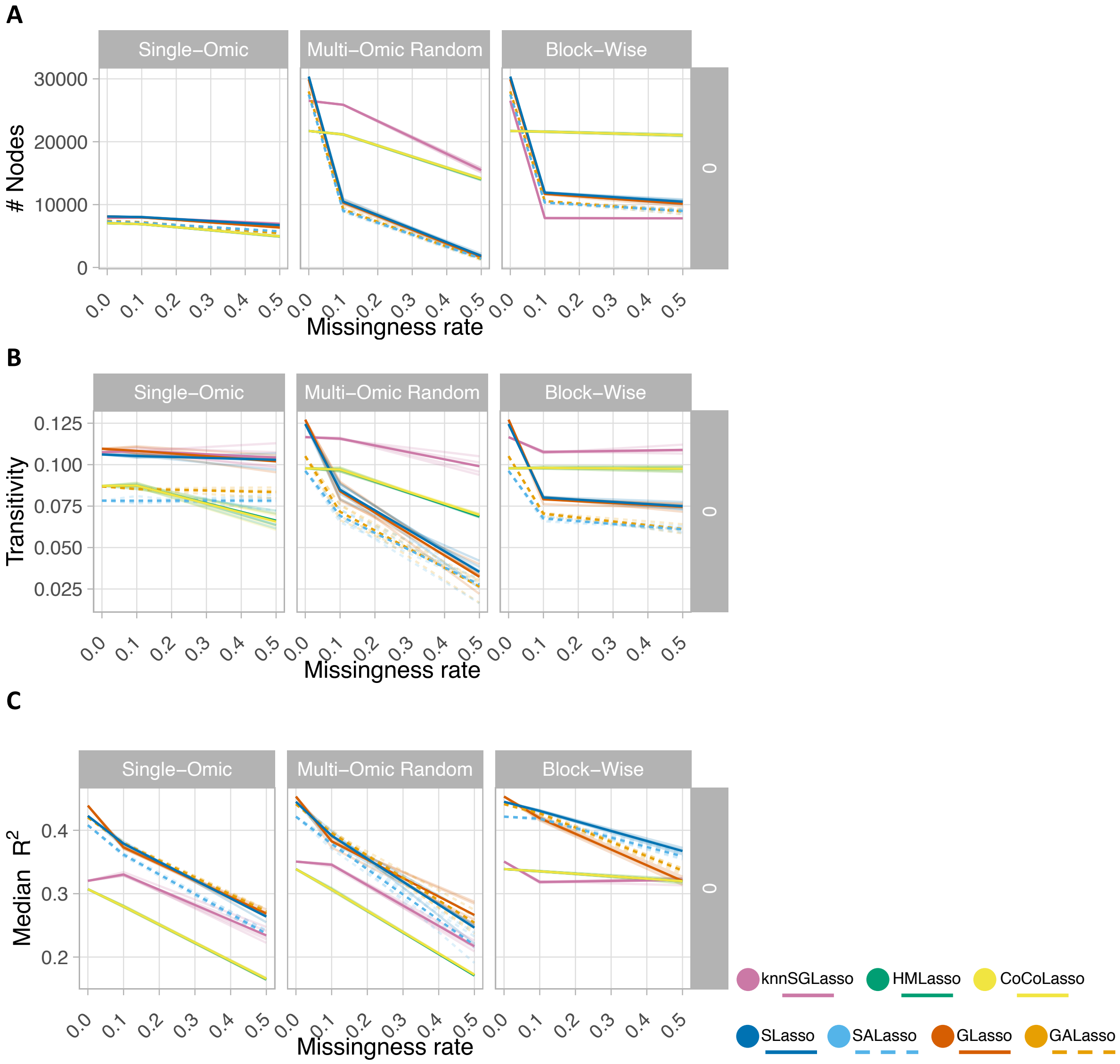


**Supplementary Figure 8.** Performance evaluation for TCGA-PRAD, **A)** Number of nodes indicating network size, **B)** Global clustering coefficient per network inferred, **C)** Median $R^2$ encompassing the model performance per network inferred.

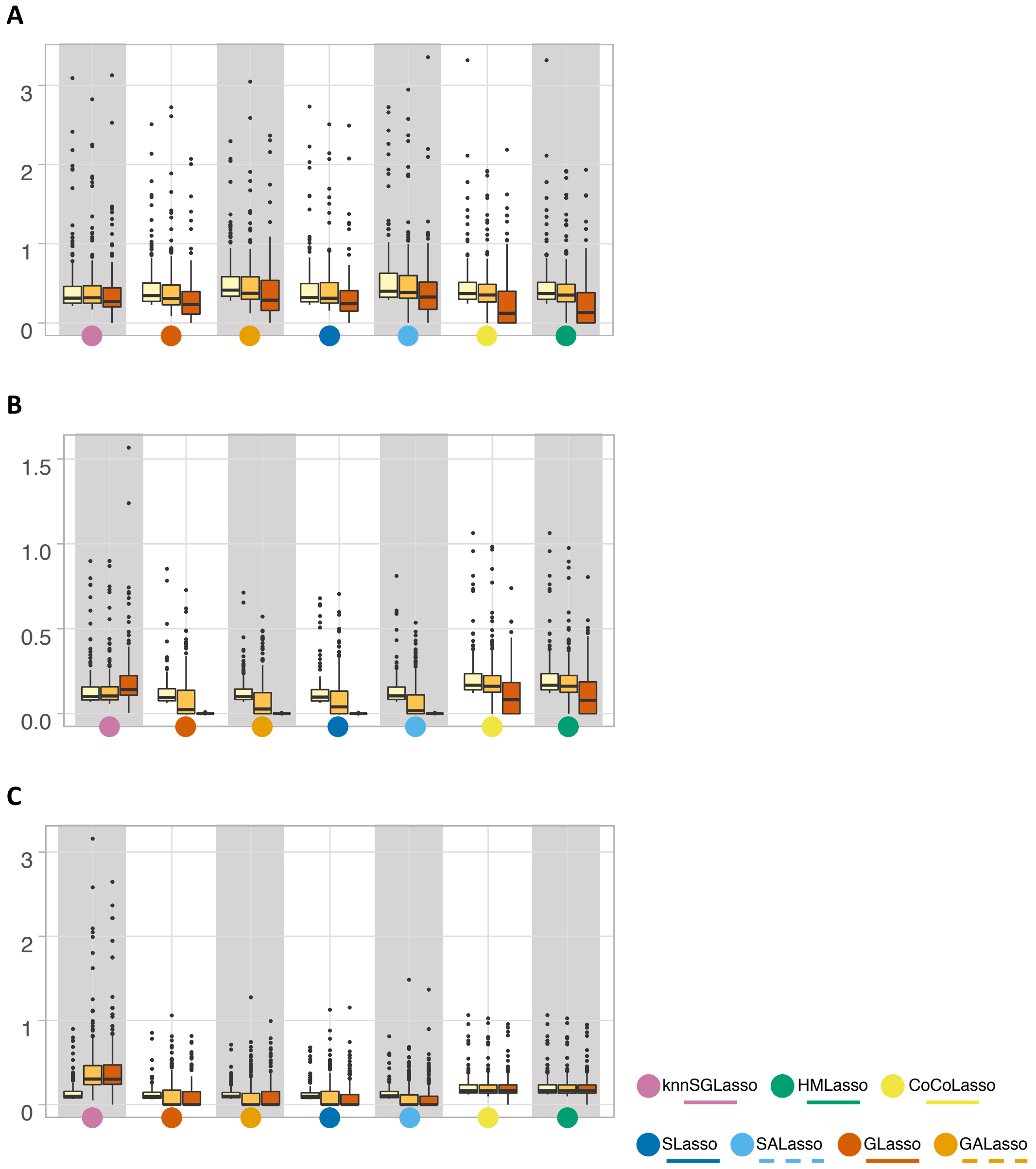


**Supplementary Figure 9.** Performance evaluation via betweenness centrality for TCGA-PRAD, **A)** Single-Omics experiment, **B)** Multi-omics experiment, **C)** Block-missingness experiment. All experiments were taken for 0%, 10%, and 50% missing ratios.

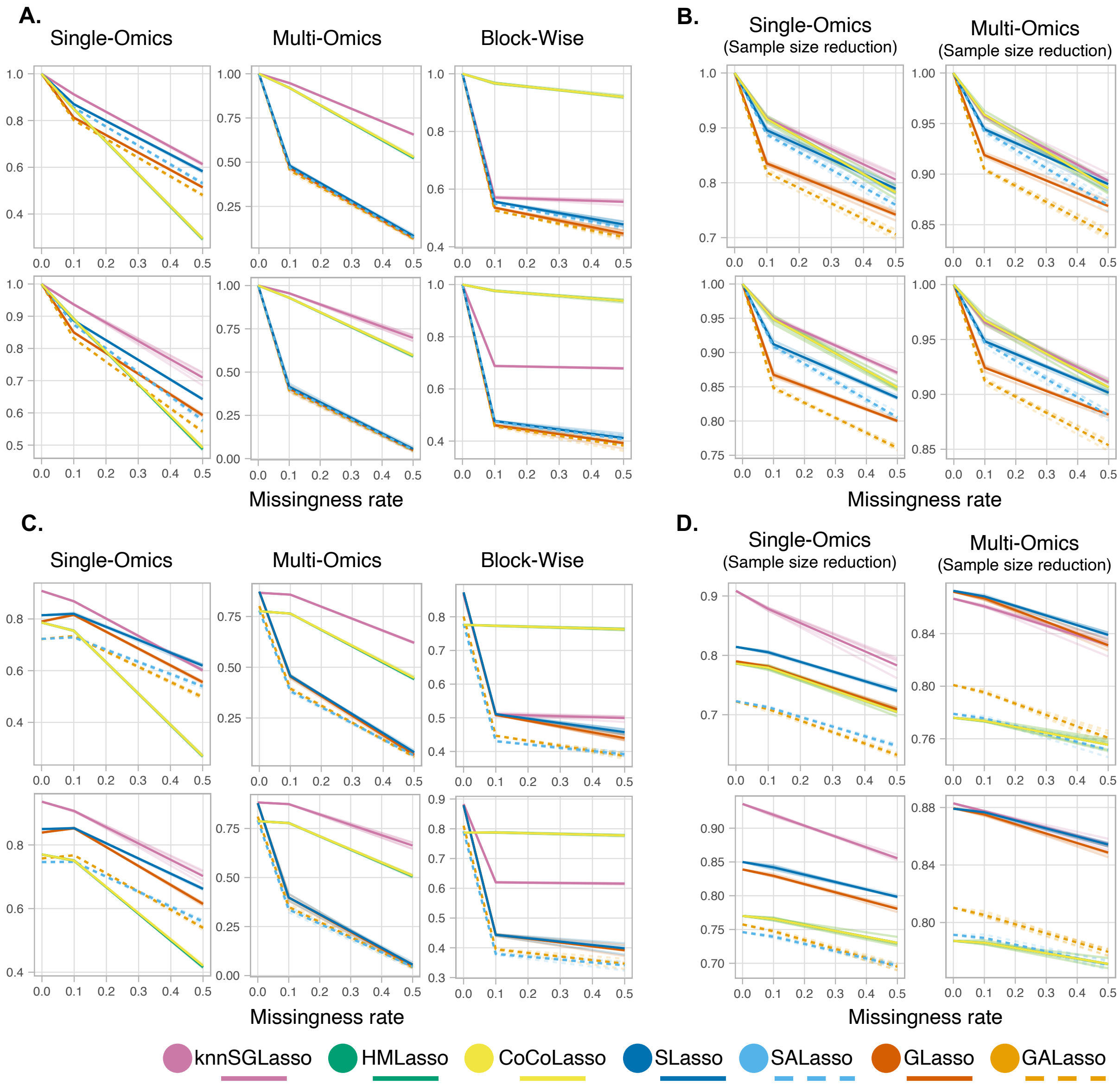


**Supplementary Figure 10.** Performance evaluation via F1-score for 10% and 50% missingness rate, **A)** & **B)** Experimental set-ups for TCGA-MIBC (top) and TCGA-PRAD (bottom) using full data as reference, **C)** & **D)** Experimental set-ups for TCGA-MIBC (top) and TCGA-PRAD (bottom) using stable networks.

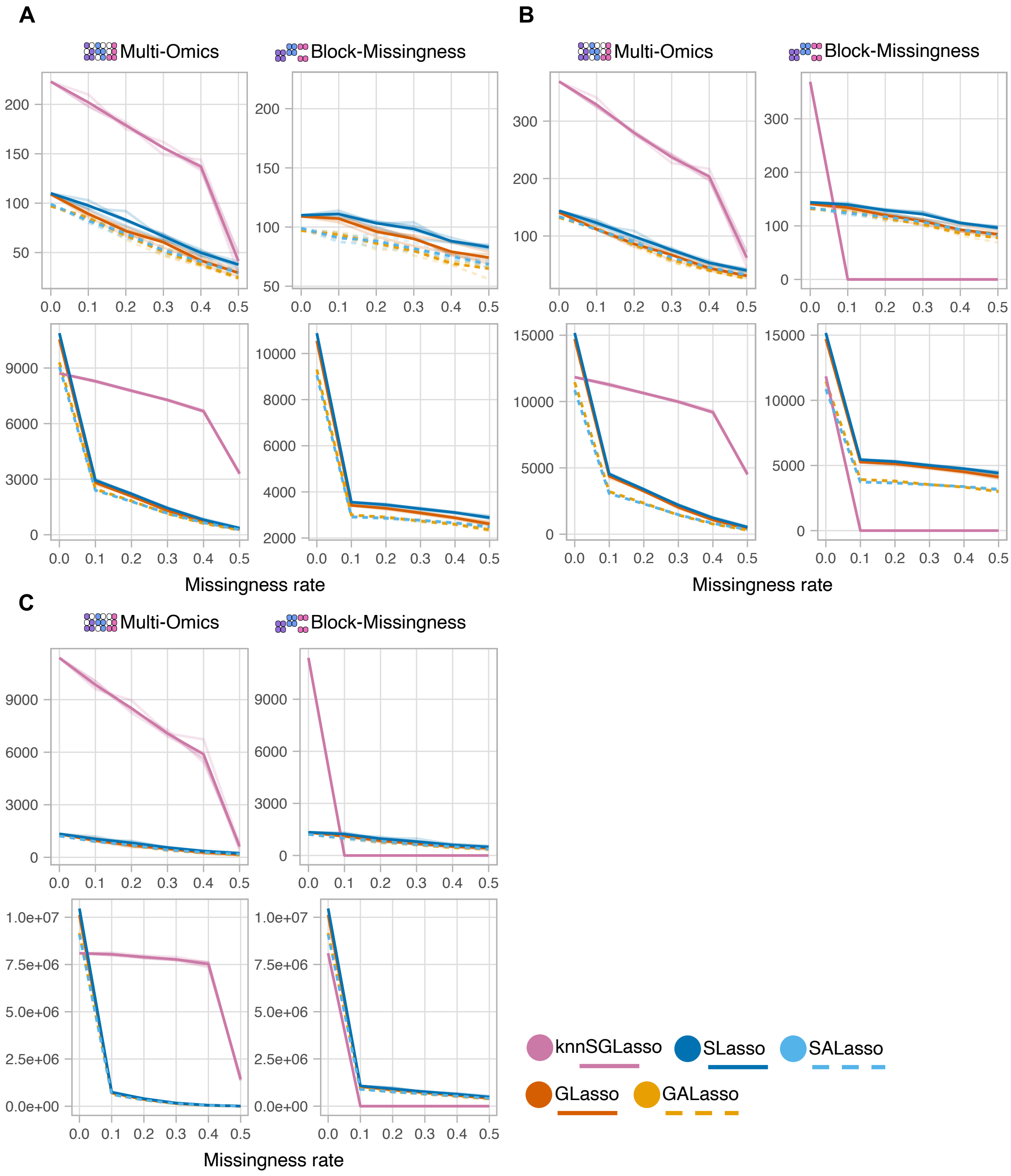


**Supplementary Figure 11.** Intra-layer evaluation via eQTM detection. **TCGA-BRCA** (top) and **TCGA-MIBC** (bottom). **A)** eQTM-linked genes detected by different inferred networks using different sparse. **B)** Number of total eQTMs detected by different inferred networks using different sparse models. **C)** Number of total eQTMs detected by Matrix eQTL based on the gene-methylation links obtained from the different inferred networks.
